# Mendelian Randomization analysis reveals a causal influence of circulating sclerostin levels on bone mineral density and fractures

**DOI:** 10.1101/455386

**Authors:** Jie Zheng, Winfried Maerz, Ingrid Gergei, Marcus Kleber, Christiane Drechsler, Christoph Wanner, Vincent Brandenburg, Sjur Reppe, Kaare M Gautvik, Carolina Medina-Gomez, Enisa Shevroja, Arthur Gilly, Young-Chan Park, George Dedoussis, Eleftheria Zeggini, Mattias Lorentzon, Petra Henning, Ulf H. Lerner, Karin Nilsson, Sofia Movérare-Skrtic, Denis Baird, Benjamin Elsworth, Louise Falk, Alix Groom, Terence D. Capellini, Elin Grundberg, Maria Nethander, Claes Ohlsson, George Davey Smith, Jonathan H. Tobias

**Affiliations:** MRC Integrative Epidemiology Unit (IEU), Bristol Medical School, University of Bristol, Oakfield House, Oakfield Grove, Bristol, BS8 2BN, United Kingdom; Clinical Institute of Medical and Chemical Laboratory Diagnostics, Medical University of Graz, Austria; SYNLAB Academy, SYNLAB Holding Deutschland GmbH, Mannheim, Germany; Vth Department of Medicine (Nephrology, Hypertensiology, Rheumatology, Endocrinology, Diabetology), Medical Faculty Mannheim, University of Heidelberg, Mannheim, Germany; Department of Internal Medicine 1, Division of Nephrology and the Comprehensive Heart Failure Center, University of Würzburg, Germany; Institute of Basic Medical Sciences, University of Oslo. Oslo, Norway; Unger-Vetlesen Institute, Lovisenberg Diaconal Hospital, Oslo, Norway; Department of Medical Biochemistry, Oslo University Hospital, Oslo, Norway; Department of Internal Medicine, Erasmus MC University Medical Center, Rotterdam, The Netherlands; Human Genetics, Wellcome Sanger Institute, Wellcome Genome Campus, Hinxton, CB10 1SA, United Kingdom; Institute of Translational Genomics, Helmholtz Zentrum München, German Research Center for Environmental Health, Neuherberg, Germany; University of Cambridge, Cambridge, UK; Department of Nutrition and Dietetics, School of Health Science and Education, Harokopio University, El. Venizelou 70, 17671, Athens, Greece; Centre for Bone and Arthritis Research, Department of Internal Medicine and Clinical Nutrition, Institute of Medicine, University of Gothenburg, Sweden; Geriatric Medicine, Institute of Medicine, University of Gothenburg, Sweden; Geriatric Medicine Clinic, Sahlgrenska University Hospital, Mölndal, Sweden; Bristol Bioresource Laboratories, Population Health Sciences, Bristol Medical School, University of Bristol, UK; Human Evolutionary Biology, Harvard University, USA; Broad Institute of MIT and Harvard, USA; Department of Human Genetics, McGill University, Quebec, Canada; Center for Pediatric Genomic Medicine, Children’s Mercy Kansas City, USA; Musculoskeletal Research Unit, University of Bristol, Level 1 Learning and Research Building, Bristol, BS10 5NB, United Kingdom

## Abstract

In bone, sclerostin is mainly osteocyte-derived and plays an important local role in adaptive responses to mechanical loading. Whether circulating levels of sclerostin also play a functional role is currently unclear, which we aimed to examine by two sample Mendelian Randomisation (MR). A genetic instrument for circulating sclerostin, derived from a genome wide association study (GWAS) meta-analysis of serum sclerostin in 10,584 European-descent individuals, was examined in relation to femoral neck bone mineral density (BMD; n= 32,744) in GEFOS, and estimated BMD by heel ultrasound (eBMD; n=426,824), and fracture risk (n=426,795), in UK Biobank. Our GWAS identified two novel serum sclerostin loci, *B4GALNT3* (standard deviation (SD)) change in sclerostin per A allele (β=0.20, P=4.6×10^−49^), and *GALNT1* (β=0.11 per G allele, P=4.4×10^−11^). *B4GALNT3* is an N-acetyl-galactosaminyltransferase, adding a terminal LacdiNAc disaccharide to target glycocoproteins, found to be predominantly expressed in kidney, whereas *GALNT1* is an enzyme causing mucin-type O-linked glycosylation. Using these two SNPs as genetic instruments, MR revealed an inverse causal relationship between serum sclerostin and femoral neck BMD (β= −0.12, 95%CI= −0.20 to −0.05) and eBMD (β= −0.12, 95%CI= −0.14 to −0.10), and a positive relationship with fracture risk (β= 0.11, 95%CI= 0.01 to 0.21). Colocalization analysis demonstrated common genetic signals within the *B4GALNT3* locus for higher sclerostin, lower eBMD, and greater *B4GALNT3* expression in arterial tissue (Probability>99%). Our findings suggest that higher sclerostin levels are causally related to lower BMD and greater fracture risk. Hence, strategies for reducing circulating sclerostin, for example by targeting glycosylation enzymes as suggested by our GWAS results, may prove valuable in treating osteoporosis.

## Introduction

Sclerostin is a glycoprotein produced by osteocytes, which is thought to play an important role in bone’s adaptive response to mechanical loading, acting within the local bone microenvironment to suppress bone formation (1). The sclerostin antibody romosozumab has recently been found to increase bone mineral density (BMD) and reduce fracture risk (2)(3), establishing sclerostin as an important drug target for osteoporosis. However, concerns of possible off-target effects were raised by the finding of an increased risk of cardiovascular events reported in a recent phase III trial of patients randomised to romosozumab or alendronate (3). Sclerostin is also detectable in the systemic circulation, the functional role of which remains unclear. Serum sclerostin exchanges with the bone micro-environment, and may simply mirror the prevalent conditions within bone. Consistent with this suggestion, serum sclerostin has been found to respond to physiological stimuli such as estrogen in parallel to alterations in local bone expression (4).

Several studies suggest that serum sclerostin exerts other roles outside the skeleton. For example, serum sclerostin has been reported to increase in chronic kidney disease (CKD), and may contribute to the pathogenesis of Chronic Kidney Disease–Mineral and Bone Disorder (CKD-MBD) (5). Serum sclerostin levels are also higher in patients with cardiovascular disease and may predict cardiovascular mortality (6). A relationship with glucose metabolism has been suggested in light by reports of higher sclerostin levels in association with type I and type II diabetes mellitus in adolescents (7) and adults (8) respectively. While these changes may represent ‘off target’ effects of sclerostin originating from bone, alternatively, sclerostin might act as an endocrine hormone, subject to regulation by as yet unidentified additional factors. Adding to this complexity, several extra skeletal tissues have been found to express sclerostin, including the liver, chondrocytes, kidney and vascular smooth muscle cells (9). Aside from extra skeletal target tissues, perturbations in serum sclerostin could conceivably affect the bone itself through consequent alterations of sclerostin levels within the local bone environment.

To the extent that circulating sclerostin levels influence bone metabolism, this may offer additional strategies for targeting sclerostin, and provide a distinct toxicity profile, compared with systemic sclerostin antibody administration producing complete abrogation of sclerostin function throughout the organism. In the present study, we aimed to establish whether circulating sclerostin exerts a causal influence on bone metabolism, using a two sample MR framework (10). With this approach, genotypic markers for the exposure of interest are first identified from a genome wide association study (GWAS), to provide an instrumental variable. Subsequently, the output of a GWAS for the outcome, obtained in a separate population, is interrogated for association with the instrumental variable for the exposure. This enables causal inferences to be drawn between exposures and outcomes based on evaluation of their genetic determinants, despite these being measured in separate populations. The recently published two sample MR, demonstrating a null association between vitamin D and fracture risk, represents a successful application of this approach (11). Therefore, we aimed to derive an instrumental variable for sclerostin from a sclerostin GWAS, and to subsequently apply this to outputs from previously published large scale bone mineral density (BMD) and fracture GWASs.

## METHODS

### Cohort Details

#### The Avon Longitudinal Study of Parents and Children (ALSPAC)

ALSPAC is a prospective birth cohort which recruited pregnant women with expected delivery dates between April 1991 and December 1992 from Bristol UK. The initial number of pregnancies enrolled is 14,541 (for these at least one questionnaire has been returned or a “Children in Focus” clinic had been attended by 19/07/1999). Of these initial pregnancies, there was a total of 14,676 foetuses, resulting in 14,062 live births and 13,988 children who were alive at 1 year of age. Detailed information on health and development of children and their parents were collected from regular clinic visits and completion of questionnaires (12)(13). Ethical approval was obtained from the ALSPAC Law and Ethics Committee and the Local Ethics Committees. Please note that the study website contains details of all the data that is available through a fully searchable data dictionary (http://www.bristol.ac.uk/alspac/researchers/our-data/).

#### Die Deutsche Diabetes Dialyse Studie (4D)

The 4D study was a prospective randomized controlled trial including patients with type 2 diabetes mellitus who had been treated by hemodialysis for less than 2 years (14). Between March 1998 and October 2002, 1255 patients were recruited in 178 dialysis centres in Germany. Patients were randomly assigned to double-blinded treatment with either 20 mg of atorvastatin (n = 619) or placebo (n = 636) once daily and were followed up until the date of death, censoring, or end of the study in March 2004. The primary end point of the 4D study was defined as a composite of death due to cardiac causes, stroke, and myocardial infarction, whichever occurred first. 4D study end points were centrally adjudicated by 3 members of the endpoints committee blinded to study treatment and according to predefined criteria. The study was approved by the medical ethics committees, and written informed consent was obtained from all participants.

#### The Gothenburg Osteoporosis and Obesity Determinants (GOOD)

The GOOD study was initiated to determine both environmental and genetic factors involved in the regulation of bone and fat mass (15). Male study subjects were randomly identified in the greater Gothenburg area in Sweden using national population registers, contacted by telephone, and invited to participate. To be enrolled in the GOOD study, subjects had to be between 18 and 20 years of age. There were no other exclusion criteria, and 49% of the study candidates agreed to participate (n = 1068). The study was approved by the ethics committee at the University of Gothenburg. Written and oral informed consent was obtained from all study participants.

#### MANOLIS Cohort

HELIC-MANOLIS (Minoan isolates) collection comprises individuals from the mountainous Mylopotamos villages, including Anogia, Zoniana, Livadia and Gonies (estimated population size of 6,000 in total) on the Greek island of Crete. It is one of the two cohorts composing the Hellenic Isolated Cohorts study (HELIC - http://www.helic.org). The specific population genetics (16) and dietary and lifestyle habits (17) of this cohort have been previously studied in the literature. The HELIC collections include blood for DNA extraction, laboratory-based haematological and biochemical measurements, and interview-based questionnaire data. The study was approved by the Harokopio University Bioethics Committee, and informed consent was obtained from human subjects.

### GWAS Meta-analysis of sclerostin

Sclerostin measures in the four cohorts were standardized to SD units. Each cohort ran a GWAS across all imputed or sequenced variants. Age, sex were included as covariates in all models, as were the first 10 principal components (PCs) to adjust for genetic stratification. Linear mixed models BOLT-LMM and GEMMA were applied to ALSPAC and MONOLIS cohort, respectively, to adjust for cryptic population structure and relatedness. The ALSPAC, 4D and GOOD cohorts were imputed using Haplotype Reference Consortium (HRC) V1.0 reference panel (MANOLIS employed whole genome sequencing). We standardized the genomic coordinates to be reported on the NCBI build 37 (hg19), and alleles on the forward strand. Summary level quality control was conducted for Europeans only in EasyQC (18). Meta-analysis (using a fixed-effects model implemented in EasyQC) was restricted to variants common to all four studies (n=5,245,208 variants), MAF >1%, and high imputation quality score (Rsq >0.8 for variants imputed in MaCH (19) and info >0.8 for variants imputed in IMPUTE (20). P < 5×10-8 in the meta-analysis was used to define genome-wide significant associations. Each locus is represented in the corresponding results table by the variant with the strongest evidence for association. A random effects model meta-analysis was also conducted using GWAMA version 2.2.2 (21). Heterogeneity was assessed using the I2 statistic and Cochrane’s Q test.

#### Conditional analysis and genetic fine mapping

To detect multiple independent association signals at each of the genome-wide significant sclerostin loci, we carried out an approximate conditional and joint genome-wide association analysis using the software package GCTA-COJO (22). SNPs with high collinearity (Correlation r^2^ > 0.9) were ignored, and those situated more than 10 Mb away were assumed to be in complete linkage equilibrium. A reference sample of 8890 unrelated individuals of ALSPAC mothers was used to model patterns of linkage desequilibrium (LD) between variants. The reference genotyping data set consisted of the same 5 million variants assessed in our GWAS. Conditionally independent variants that reached GWAS significance were annotated to the physically closest gene with the Hg19 Gene range list available in dbSNP (https://www.ncbi.nlm.nih.gov/SNP/).

#### Estimation of SNP heritability using LD-score regression

To estimate the amount of genomic inflation in the data due to residual population stratification, cryptic relatedness and other latent sources of bias, we used LD score regression (23). LD scores were calculated for all high-quality SNPs (i.e., INFO score > 0.9 and MAF > 0.1%) from the meta-analysis. We further quantified the overall SNP-based heritability’s with LD score regression using a subset of 1.2 million HapMap SNPs.

#### Estimation of genetic correlations using LD Hub

To estimate the genetic correlation between sclerostin and bone phenotypes related to osteoporosis, we used a recent method based on LD score regression as implemented in the online web utility LD Hub (24). This method uses the cross-products of summary test statistics from two GWASs and regresses them against a measure of how much variation each SNP tags (its LD score). Variants with high LD scores are more likely to contain more true signals and thus provide a greater chance of overlap with genuine signals between GWASs. The LD score regression method uses summary statistics from the GWAS metaanalysis of sclerostin and the bone phenotypes, calculates the cross-product of test statistics at each SNP, and then regresses the cross-product on the LD score.

#### Genetic colocalization analysis between sclerostin and bone phenotypes

We used a stringent Bayesian model (coloc) to estimate the posterior probability (PP) of each genomic locus containing a single variant affecting both serum sclerostin level and bone phenotypes (25). A lack of evidence (i.e. PP < 80%) in this analysis would suggest that one of the causal variants for protein is simply in LD with the putative causal variant for the trait (thus introducing genomic confounding into the association between sclerostin level and bone phenotypes). We treated colocalised findings (PP>=80%) as “Colocalised”, and other findings that did not pass colocalization as “Not colocalised”.

### Mendelian randomization

#### Sclerostin versus bone phenotypes

We undertook two sample MR (26) to evaluate evidence of a causal relationship between sclerostin measured in plasma and bone phenotypes. We looked up GWAS results in distinct cohorts to those used for the sclerostin GWAS, namely GEFOS in the case of femoral neck and lumbar spine BMD (N=32,961) (27), and UK Biobank in the case of eBMD (N=426,824) (28), and (self-reported) fracture risk (N=426,795) (29). In this initial analysis, sclerostin was treated as the exposure and bone traits as the outcomes, using sclerostin associated SNPs as the instrumental variables. The multiple testing p-value threshold was calculated as p = 0.05 divided by the derived number of independent tests. We used the random effect inverse variance weighted (IVW) (30) and Wald ratio (31) methods to obtain MR effect estimates. Heterogeneity analysis of the instruments were conducted using Cochran Q test. Results were plotted as forest plots using code derived from the ggplot2 package in R (32). All MR analysis were conducted using the MR-Base TwoSampleMR R package (github.com/MRCIEU/TwoSampleMR) (33). To identify potential pleiotropic pathways, we conducted a phenome-wide association study (PheWAS) of the two top hits, rs215226 and rs7241221 using the MR-Base PheWAS tool (http://phewas.mrbase.org/) (33), applying a Bonferroni P value threshold of 3×10^−6^, to account for multiple testing (number of tests=22027).

#### Bidirectional MR

To investigate the possibility that BMD causally affects levels of serum sclerostin, we used summary results data from 49 conditionally independent autosomal variants reported in a BMD GWAS using 32,744 GEFOS individuals (27). We looked up these variants in summary results GWAS data on sclerostin from our meta-analysis and found 39 instruments (including two proxy SNPs) for femoral neck BMD and 38 instruments (including two proxy SNPs) for lumbar spine BMD. When applying bidirectional MR, we assume that SNPs used to proxy BMD exert their primary association on BMD, and that any correlation with sclerostin levels is a consequence of a causal effect of BMD on sclerostin. We therefore applied Steiger filtering (34) and included 33 femoral neck BMD SNPs and 35 lumbar spine BMD SNPs which exert their primary effect on BMD. See supplementary methods for additional details.

### Functional annotation

#### Predicted regulatory elements of the top association signals

For each locus associated with sclerostin, we identified all SNPs with high LD from the top signal (LD r^2^>0.8) and identify their DNA features and regulatory elements in non-coding regions of the human genome using Regulomedb v1.1 (http://www.regulomedb.org/) (35).

#### ATAC-seq lookup in the top association regions

The Assay for Transposase Accessible Chromatin with high-throughput sequencing (ATAC-seq) is a method for mapping chromatin accessibility genome-wide (36). We intersected the sclerostin associated SNPs (or proxy SNPs with LD r^2^ > 0.8) with ATAC-seq data generated from the proximal and distal femur of an E15.5 mouse (37)(36) lifted over to the orthologous positions in human genome build hg19).

#### Gene expression quantitative trait loci (eQTLs) lookups for sclerostin signals

We investigated whether the SNPs influencing serum sclerostin level were driven by cis-acting effects on transcription by evaluating the overlap between the sclerostin associated SNPs and eQTLs within 500kb of the gene identified, using data derived from all tissue types from the GTEx consortium v7 (38). Evidence of eQTL association was defined as P < 1×10-4 and evidence of overlap of signal was defined as high LD (r^2^ ≥ 0.8) between eQTL and sclerostin associated SNPs in the region. Where eQTLs overlapped with sclerostin associated SNPs, we used the colocalization analysis to estimate the posterior probability (PP) of each genomic locus containing a single variant affecting both serum sclerostin level and gene expression level in different tissues. Equivalent analyses were performed in human primary osteoblast cells (39), and cells derived from iliac crest bone from 78 postmenopausal women (40) (see supplementary methods for further details).

#### Methylation quantitative trait loci (mQTLs) lookups for sclerostin signals

We also investigated whether the SNPs influencing serum sclerostin level were mediated by cis-acting effects on DNA methylation by evaluating the overlap between the sclerostin associated SNPs and mQTLs within 500kb of the gene identified, using data measured in blood cells. We queried data from mQTLdb, which contained mQTL information at 5 different time points, ALSPAC children at Birth, Childhood and Adolescence and ALSPAC mothers during Pregnancy and at Middle age. This data is from the ARIES resource (41). Evidence of mQTL association was defined as P < 1×10^−4^ and evidence of overlap of signal was defined as high LD (r^2^ ≥ 0.8) between mQTL and sclerostin associated SNPs in the region.

## RESULTS

### Genome-wide association study meta-analysis of serum sclerostin

GWAS results were available in 10,584 participants across four cohorts (see Table 1). Supplementary Figures 1 and 2 show the Manhattan plot and QQ plot of association results from the fixed-effect meta-analysis of sclerostin in Europeans, respectively (see Supplementary Figure 3 for the QQ plots for each study). There was little evidence of inflation of the association results as the genomic inflation factor *λ* was 1.051 and the LD score regression intercept 1.015 (23). Therefore, genomic control correction was not applied to any of the meta-analysis results. LD score regression revealed that all common variants we included in the meta-analysis explained 16.3% of the phenotypic variance of sclerostin (H^2^=0.163, S.E.=0.052, P=0.0017).

**Table 1.** Study information of the cohorts involved in the sclerostin GWAS meta-analysis.

| COHORT | N<br>SCLEROSTIN | N FEMALE<br>SCLEROSTIN | N<br>GWAS | AGE (SD) | ETHNIC OR<br>RACIAL<br>GROUP | FAMILY<br>STRUCTURE | FAMILY<br>STRUCTURE<br>ADJUSTMENT | ASSAY DETAILS |
| --- | --- | --- | --- | --- | --- | --- | --- | --- |
| ALSPAC | 8772 | 6371 | 7292 | Children:<br>9.9 (0.3)<br>Mothers:<br>48 (4.4) | European | Mothers and<br>Children | BOLT-LMM | Biomedica<br>Human<br>Sclerostin<br>ELISA |
| 4D | 1041 | 468 | 1041 | 66.3 (8.36) | European | Unrelated | No adjustment | TECO |
| MANOLIS | 1316 | 729 | 1316 | 61 (19) | European | Isolated<br>population | GEMMA | OLINK |
| GOOD | 935 | 0 | 935 | 18.9(0.56) | European | Unrelated | No adjustment | TECO |

ALSPAC and GOOD represent general population cohorts, MANOLIS is an isolated population from Crete, and 4D is a cohort of patients receiving haemodialysis for end-stage renal failure.

Two loci were identified to be associated with serum sclerostin at genome-wide significance level. 69 SNPs in the *B4GLANT3* gene region were associated with sclerostin level (Figure 1A). The top hit rs215226 at this locus (β = 0.205 SD change in serum sclerostin per A allele, SE = 0.014, P = 4.60 x 10^−49^, variance explained = 1.99%) was previously reported to be associated with BMD estimated by heel ultrasound (eBMD) (28). The second locus was in the *GALNT1* gene region (Figure 1B) with the top hit rs7241221 (β = 0.109 SD per G allele, SE = 0.017, P = 4.40 x 10^−11^, variance explained = 0.39%). In addition, one SNP near the *TNFRSF11B* region encoding OPG, rs1485303, showed a suggestive association with sclerostin (β = 0.074 SD per G allele, SE = 0.014, P = 7.70 × 10^−8^, variance explained = 0.26%) (Figure 1C). The latter SNP is in moderate LD with a previously reported BMD associated SNP rs1353171 (r^2^=0.277 in the 1000 Genome Europeans; distance between the two SNPs= 103,258 base pairs) (42). The association signals of the GWAS meta-analysis can be found in Table 2 and Supplementary Table 1. Results of the random effects meta-analysis were identical to those of the fixed effects meta-analysis (results not shown). The degree of heterogeneity was low across studies for the *GALNT1* signal (I^2^=0) and *TNFRSF11B* signal (I^2^=0.53) but relatively high for *B4GALNT3* signal (I^2^=0.78) (Supplementary Table 2). The latter appeared to reflect a relatively strong association in 4D, suggesting as possible interaction with the presence of CKD. Conditional analyses on the lead SNP in each association locus yielded no additional independent signals reaching genome-wide significance. The association of the two sclerostin SNPs with femoral neck, lumbar spine and heel BMD were presented in Supplementary Table 3.

**Figure 1.**
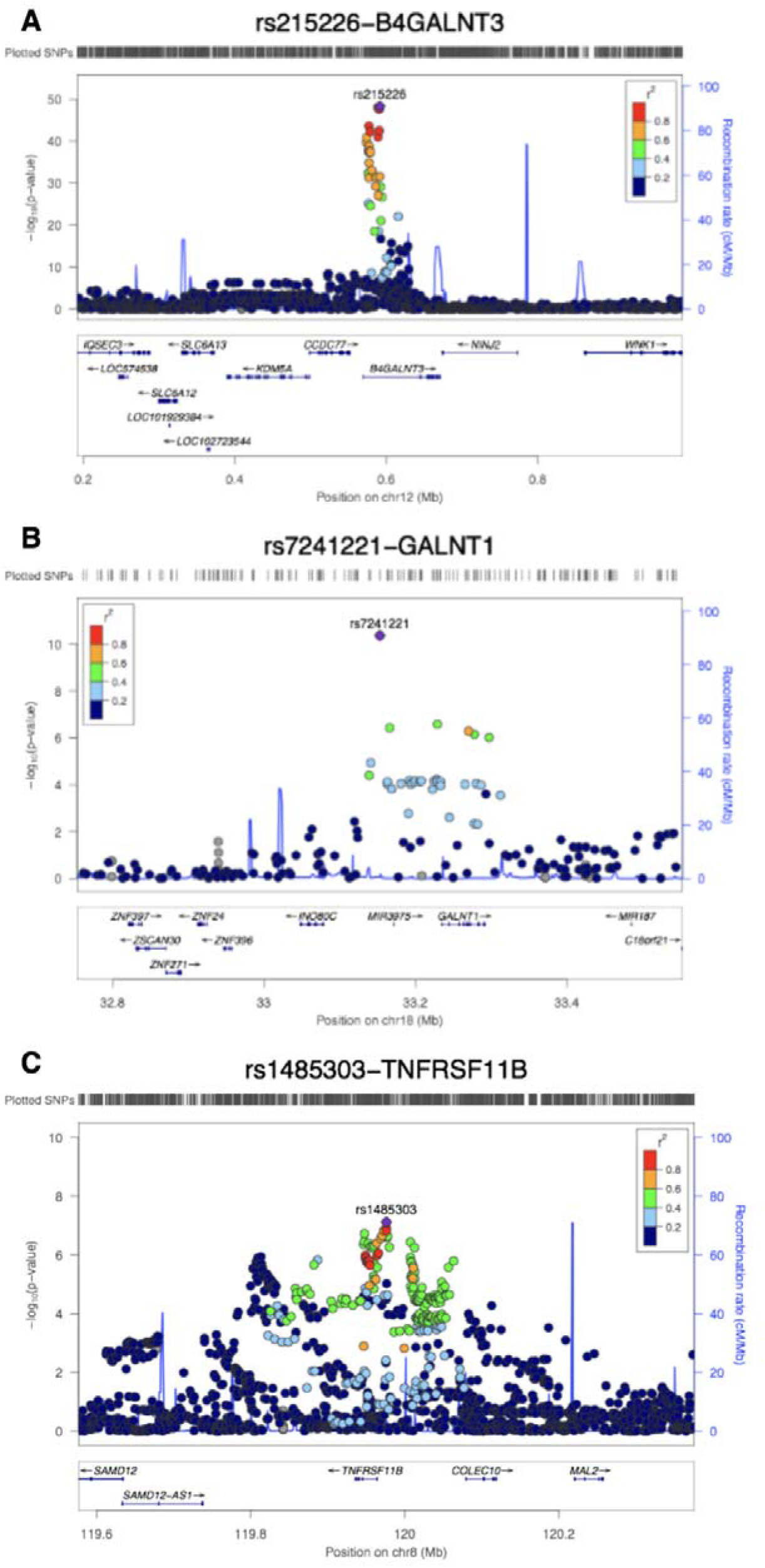
Regional association plots and ENCODE annotation of the loci that reached or marginally reached genome-wide significance (P < 5 × 10^−8^) in the meta-analysis. The X-axis indicates the physical position of each SNP on the chromosome specified, whereas the Y-axis denotes the evidence of association shown as −log(P-value). A) the *B4GLANT3* region; 2) the *GALNT1* region and 3) the *TNFRSF11B* region.

**Table 2.** Meta-analysis results for loci that reached or marginally reached genome-wide significance (P < 5 × 10 ^8^).

| Locus | SNP | EA | OA | EAf | GENE | BETA | SE | P | Q | Q_P | I<sup>2</sup> |
| --- | --- | --- | --- | --- | --- | --- | --- | --- | --- | --- | --- |
| CHR12:591300 | rs215226 | A | G | 0.597 | *B4GALNT3* | 0.205 | 0.014 | $4.60 \times 10^{-49}$ | 13.43 | 0.0038 | 0.777 |
| CHR18:33152792 | rs7241221 | G | A | 0.772 | *GALNT1* | 0.109 | 0.017 | $4.40 \times 10^{-11}$ | 0.955 | 0.812 | 0 |
| CHR8:119976256 | rs1485303 | G | A | 0.568 | *TNFRSF11B* | 0.074 | 0.014 | $7.70 \times 10^{-08}$ | 6.416 | 0.093 | 0.532 |

Note: Locus (chromosome and position of the SNP), EA (effect allele), OA (other allele), EAF (effect allele frequency), GENE (nearest gene to the sclerostin associated SNP); BETA (SD change in serum sclerostin per effect allele), SE (standard error) and P (p-value)) Heterogeneity test (Q (Cochrane's Q statistics), Q\_P (Cochrane's Q P value), I<sup>2</sup> (I<sup>2</sup> statistics))

Only minor genetic influences on serum sclerostin were observed within the *SOST* gene (Chromosome 17, 41731099 to 41936156), which produces sclerostin. In total, four *SOST* SNPs have previously been reported in association with BMD (28)(43), and one with fracture (11). However, proxy SNPs for these SNPs showed little or no association with serum sclerostin (Supplementary Table 4). We used the GWAS results to estimate the genetic correlation between sclerostin and 11 bone phenotypes using LD score regression via LD Hub (24). We observed no strong evidence of genetic correlation between sclerostin SNPs and the bone phenotypes we tested. (Supplementary Table 5).

### Mendelian randomization and colocalization analysis of sclerostin and bone phenotypes

Using two sample Mendelian Randomization (MR), we applied our GWAS results to larger datasets in order to examine putative causal relationship between sclerostin and bone phenotypes. The two SNPs robustly associated with sclerostin (rs215226 in *B4GALNT3* region and rs7241221 in *GALNT1* region) were used as genetic instruments of the exposure. DXA-derived BMD and eBMD results from 32,961 GEFOS individuals (27) and 426,824 UK Biobank individuals (28) respectively, and self-reported fracture of 426,795 individuals from UK Biobank, were used as outcomes. Contrasting with the genetic correlation results, based on inverse variance weighted (IVW) analysis, we observed that a higher level of serum sclerostin was causally related to lower femoral neck BMD (β= −0.123 SD change in BMD per SD increase in sclerostin, 95%CI= −0.195 to −0.051, P=0.00074), lower eBMD (β= −0.122 SD change in BMD per SD increase in sclerostin, 95%CI= −0.140 to −0.104, P= 1.29×10^−38^) and higher fracture risk (β=1.117 odds ratio (OR) of fracture per SD increase in sclerostin, 95%CI= 1.009 to 1.237, P= 0.034) (Figure 2). Point estimates for the association between sclerostin level and lumbar spine BMD were similar to those observed for femoral neck and eBMD, however 95% confidence limits were wide and crossed zero. MR analyses based on single SNPs suggested that both SNPs contributed to the overall causal effects we observed (Supplementary Table 6). Heterogeneity analysis of the instruments suggested that causal estimates were consistent across both SNPs (P values of Cochran Q test > 0.05), with the exception of estimates for lumbar spine BMD and fracture risk, likely reflecting the lower power in these instances (44). Genetic signals for sclerostin and eBMD colocalized at both the *B4GALNT3* locus (PP=99.7%) and *GALNT1* locus (PP=99.8%) (Supplementary Table 7), further strengthening the evidence of the putative causal relationship between sclerostin and eBMD.

**Figure 2.**
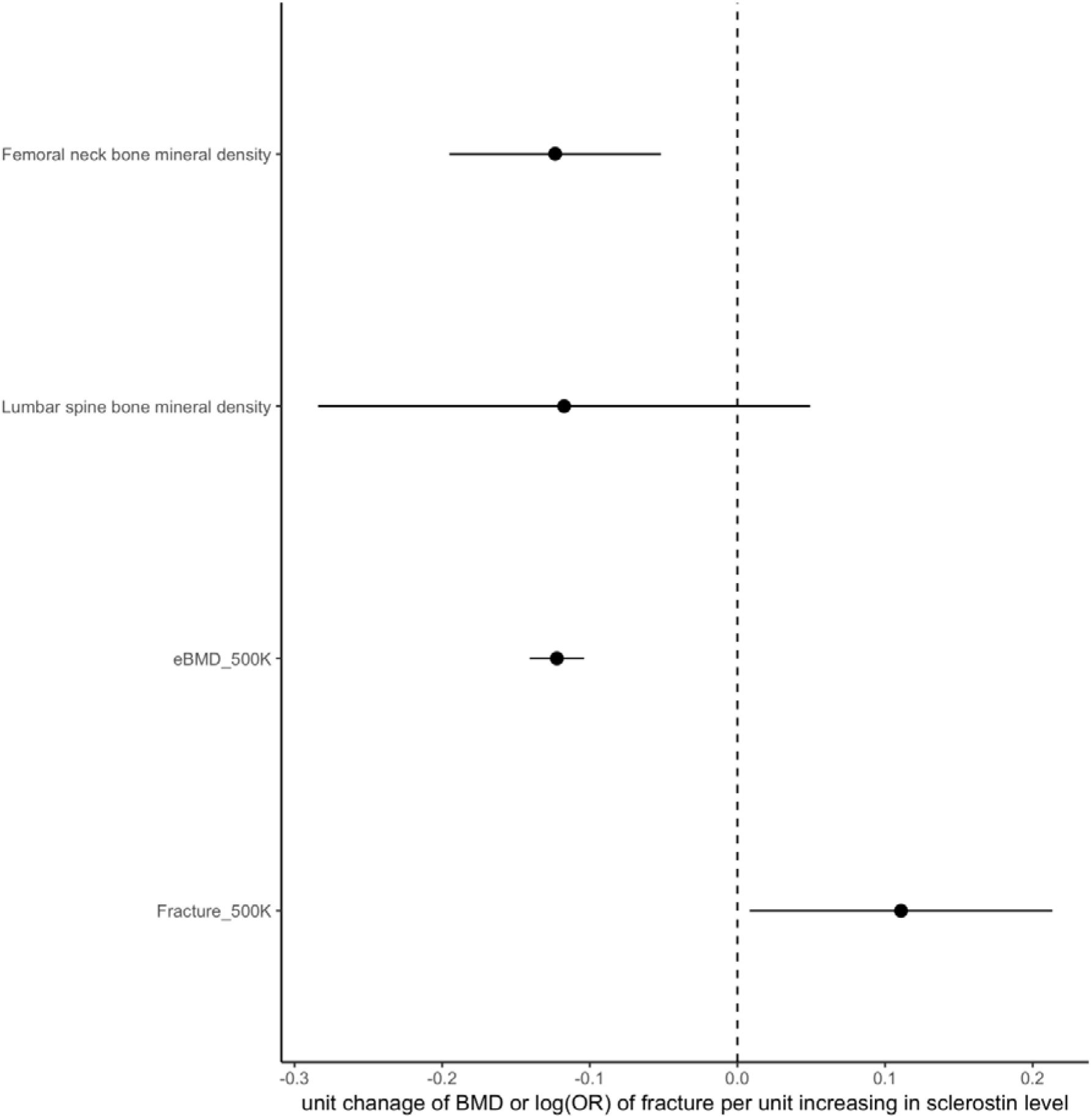
Forest plot of putative causal relationship between serum sclerostin and bone phenotypes using Mendelian randomization. X-axis represents the causal estimates and 95% confidence intervals of SD change in BMD/eBMD and OR for fracture, per SD increase in sclerostin, as calculated by inverse variance weighted method. Y-axis lists the four bone phenotypes used in the MR analysis.

Since our genetic instrument for sclerostin was only based on two SNPs, we were unable to apply methods such as Egger regression to test the assumption that this instrument affects BMD via sclerostin as opposed to a separate causal pathway (i.e. horizontal pleiotropy). Therefore, to evaluate the possible role of pleiotropy, we performed a phenome-wide association study of sclerostin associated signals. rs215226 and rs1485303 were associated with other heel ultrasound parameters closely related to eBMD; rs215226 was also associated with height and body mass, based on Bonferroni P value threshold of 3×10^−6^ (Supplementary Table 8 and 9). The strength of the association between rs215226 and height, which approached that for eBMD, raised the possibility of pleiotropy. However, in subsequent analyses, the relation between observed height and eBMD in UK Biobank was found to be very weak, as reflected by Pearson correlation (r) <0.1 (Supplementary Table 10). Moreover, associations between the three top hits and sclerostin were unchanged following additional adjustment for height, as assessed in ALSPAC (Supplementary Table 11). Taken together, these findings suggest that a separate pathway via height is unlikely to contribute to the relationship we observed between our genetic instrument for sclerostin and BMD.

We also undertook bidirectional MR (45) to evaluate the possibility of reverse causality between femoral neck BMD (or lumbar spine BMD) and serum sclerostin level, based on SNPs identified in GEFOS (27). We modelled femoral neck BMD and lumbar spine BMD as our exposure and serum sclerostin level as our outcome. Having used Steiger filtering to identify SNPs predominantly related to BMD as opposed to sclerostin (34), 33 of the 39 femoral neck BMD SNPs and 35 of the 38 lumbar spine BMD SNPs were found to exert their primary effect on BMD (Supplementary Table 12). IVW results using 33 femoral neck BMD SNPs suggested a positive relationship between femoral neck BMD and sclerostin (β=0.167 SD change in sclerostin per SD change in BMD, 95%CI= 0.050 to 0.285, P=0.005) while IVW results using 35 lumbar spine BMD SNPs suggested a positive relationship between lumbar spine BMD and sclerostin (β=0.195 SD change in sclerostin per SD change in BMD, 95%CI=0.080 to 0.310, P=0.0009).

Sensitivity analyses were performed to evaluate horizontal pleiotropy. MR Egger and Weighted Median methods showed similar causal effects (Supplementary Table 13 and Supplementary Figure 4A, 4B, 4C and 4D). There was no strong evidence of horizontal pleiotropy for femoral neck BMD (Egger regression intercept = −0.0086, P = 0.369) or lumbar spine BMD (Egger regression intercept = −0.0104, P = 0.370). The heterogeneity test suggested some heterogeneity across instruments for femoral neck (Cochrane Q = 36.284, P = 0.0285) and lumbar spine BMD (Cochrane Q = 54.721, P = 0.0136). To address the heterogeneity across instruments, we further detected 3 outlier SNPs for femoral neck BMD and lumbar spine BMD separately using Cochrane test implemented in “RadialMR” R package (46)(47). The radial plot can be found in Supplementary Figure 5. As a sensitivity analysis, we removed the outlier SNPs and IVW results using the remaining 30 femoral neck BMD and 32 lumbar spine BMD SNPs still suggested a positive relationship between femoral neck BMD and sclerostin (β=0.149, 95%CI=0.045 to 0.253, P=0.005) as well as lumbar spine BMD and sclerostin (β=0.229, 95%CI=0.126 to 0.332, P=1.4×10^−5^) (Supplementary Table 13).

### Functional follow-up

#### Predicted regulatory elements of the top association signals

We used Regulomedb to identify SNPs in high LD with the top association signals (r^2^>0.8), which are likely be located within DNA regulatory elements (35). Among the 24 tested SNPs, we found three proxy SNPs showing high Regulomedb score (Supplementary Table 14): rs1872426 (r^2^=0.87 with the leading SNP rs1485303 in the *TNFRSF11B* region) had a score of 1f (likely to comprise a DNA regulatory element and linked to expression of a gene target); rs215224 and rs4980826 (r^2^=1 and 0.85 with the leading SNP rs215226 respectively in the *B4GALNT3* region) had a score of 2b (likely to comprise a DNA regulatory element) (see Supplementary Figure 6 for rs215224). We also examined whether variants we identified are located at sites of accessible chromatin, indicative of DNA transcriptional activator binding, by interrogating ATAC-seq data (37) (36) generated from the proximal and distal femur (48). In the *TNFRSF11B* locus, one SNP, rs2073618, overlapped with an ATAC-seq peak, while in the *B4GALNT3* locus, four SNPs (rs12318530, rs215223, rs215224 and rs215225) fell in the same ATAC-Seq and Chip-Seq peak (Supplementary Figure 7 and Supplementary Table 15).

#### Expression QTLs lookups for sclerostin signals

To evaluate if the sclerostin association signals influence transcription of neighbouring genes, we cross-referenced the sclerostin SNPs with *cis*-expression data in tissues measured in the GTEx consortium v7 (38), primary osteoblast cell lines (39) and iliac crest bone biopsies (40). The top association signal within the *B4GALNT3* region, rs215226, showed a strong positive association with *B4GALNT3* gene expression in arterial and ovarian tissue (Supplementary Table 16). In contrast, a weaker inverse association with *B4GALNT3* expression was observed in nervous tissue. Colocalization analysis yielded strong evidence that sclerostin levels share the same causal variant at this locus with arterial and ovarian *B4GLANT3* expression (probability > 99%), whereas there was no evidence of co-localisation in the case of nervous tissue (Supplementary Table 17). The association signal in the *GALNT1* region was positively associated with *GALNT1* expression in adipose tissue, and that of the neighbouring gene, *INO80C*, in adipose and heart tissue. In contrast, sclerostin association signals in *B4GALNT3, TNFRSF11B* and *GALNT1* were not associated with *cis*-regulation of mRNA expression in osteoblast cells (Supplementary Table 18) or iliac crest bone biopsies (Supplementary Table 19). In *trans*-eQTL analyses, the *GALNT1* signal was related to lower *SOST* mRNA levels in iliac crest bone biopsies, particularly in Affymetrix chip analyses, whereas no association was observed for *B4GALNT3* or *TNFRSF11B* (Supplementary Table 20).

#### Methylation QTLs lookups for sclerostin signals

To evaluate if our top association signals have the potential to influence DNA methylation, we cross-referenced the sclerostin SNPs with methylation data as measured at adjacent CpG sites in blood cells obtained at five different time points (ALSPAC children at Birth, Childhood and Adolescence; ALSPAC mothers during Pregnancy and at Middle age (41)). We found that our top hit rs215226, within *B4GALNT3*, was consistently associated with extent of methylation at cg20907806 and cg26388816 across four time points (Supplementary Table 21). These two CpG sites are located in a CpG island that overlapped with one of the four ATAC-seq peaks within the *B4GALNT3* region (Supplementary Figure 7, upper plot). The other potential signal near *TNFRSF11B*, rs1485303, was associated with methylation level at cg13268132 and cg17171407 at all five time points.

#### *B4galnt3* mRNA expression pattern (see supplementary methods)

In murine gene expression studies, *B4galnt3* mRNA was expressed at highest levels in kidney, with relatively high levels of expression also observed in bone, particularly cortical bone (Figure 3A). *B4galnt3* mRNA was subsequently found to be expressed at relatively high levels in osteoblast cultures derived from neonatal mouse calvariae. The expression in osteoblast cultures was similar after 2 and 4 days of culture in osteogenic media (Figure 3B). After 7 days culture there was a suggestive increase in *B4galnt3* expression. No expression was detected in *in vitro* cultured mouse bone marrow macrophages or osteoclasts (Figure 3B).

**Figure 3.**
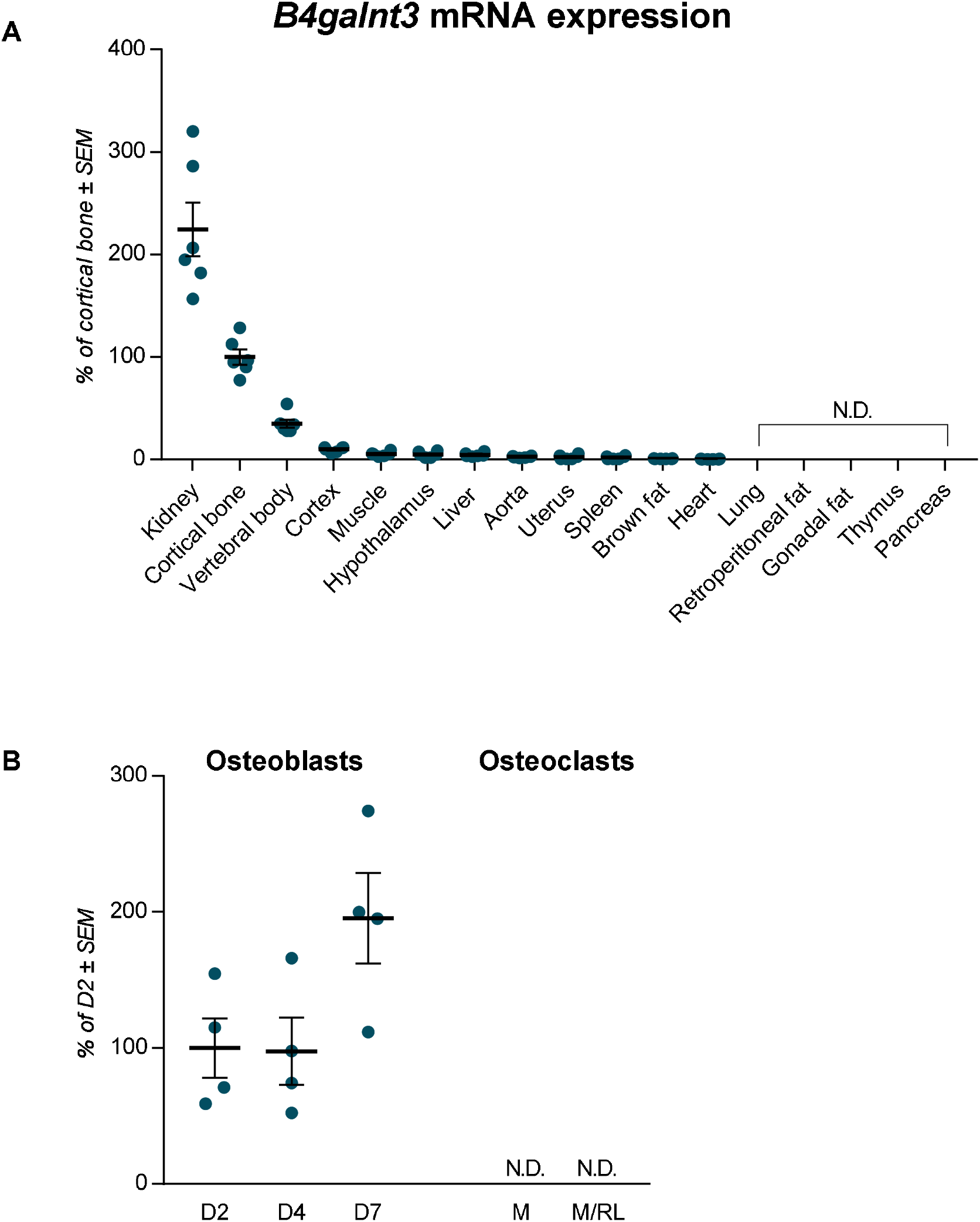
*B4galnt3* expression in mice and in *in vitro* cultured osteoblasts and osteoclasts. A. *B4galnt3* mRNA tissue expression in adult female mice, presented as % of cortical bone ±SEM (n=6). B. *B4galnt3* mRNA expression in mouse calvarial osteoblasts cultured in osteogenic media for 2, 4 and 7 days (D). Mouse bone marrow macrophages cultured with M-CSF (M), or M-CSF+RANKL (M/RL) to induce osteoclasts differentiation. % of expression D2 ±SEM. ND = not detectable

## DISCUSSION

In this two sample MR study, we first performed a GWAS meta-analysis of serum sclerostin, identifying two genome-wide significant loci (*B4GALNT3* and *GALNT1*), and a further locus close to genome wide significance (*TNFRSF11B*). Together, common genetic variants explained 16% of the variance of serum sclerostin. By using rs215226 in *B4GALNT3* and rs7241221 in *GALNT1* as instrumental variables, we subsequently examined causal relationships between serum sclerostin and BMD, finding evidence of an inverse relationship between sclerostin and BMD, and a positive relationship with fracture risk. Randomized control trial findings demonstrate that systemic administration of the sclerostin inhibitor, romosozumab, increases BMD and reduces fracture risk (2)(3). Our present findings suggest that a similar benefit may be obtained by strategies aiming to reduce levels of circulating sclerostin. However, whereas complete abrogation of sclerostin function throughout the organism following romosozumab administration appears to cause off target effects such as cardiovascular toxicity, pharmacological reduction of circulating sclerostin levels could conceivably have a distinct safety profile.

As well as providing evidence for a causal effect of circulating sclerostin on BMD and fracture risk, our GWAS findings raise the intriguing possibility that circulating sclerostin levels are influenced by the activity of glycosylation enzymes, suggesting a means by which these levels might be targeted pharmacologically. *B4GALNT3* is an N-acetyl-galactosaminyltransferase enzyme which adds a terminal LacdiNAc disaccharide to target glycocoproteins, while *GALNT1* is an enzyme causing mucin-type O-linked glycosylation. Association signals with sclerostin at these two loci colocalized with those for eBMD, consistent with the possibility that genetic variation in *B4GALNT3* and *GALNT1* alters BMD via changes in serum sclerostin.

*B4GALNT3* has previously been reported to be expressed in multiple human tissues, including arterial, gastrointestinal tract and testis (49). In the present study, in mice, *B4GALNT3* mRNA was expressed at highest levels in kidney and cortical bone. This finding is consistent with high levels of *B4GALNT3* mRNA expression in renal tissue from human fetal samples in the NIH Roadmap Epigenomics Mapping Consortium (see http://www.roadmapepigenomics.org/data/). The *B4GALNT3* locus we identified showed strong *cis*-regulatory activity in eQTL studies in arterial tissue, which was confirmed by the colocalization analysis. The effect allele associated with higher sclerostin levels was associated with considerably greater *B4GALNT3* mRNA levels in arterial tissue, suggesting that B4GALNT3 acts to increase sclerostin levels.

Sclerostin is a glycoprotein which, like *B4GALNT3*, is also expressed at a number of sites outside the skeleton including arterial tissue (50). Whereas sclerostin protein is detected in multiple tissues, extra-skeletal sclerostin mRNA is mainly expressed in the kidney, reflecting renal tubular expression (www.proteinatlas.org), suggesting this may be the predominant extra-skeletal source of circulating sclerostin. It’s tempting to speculate that sclerostin acts as a substrate for *B4GALNT3*, such that formation of a terminal LacdiNAc moiety protects sclerostin from degradation and clearance from the circulation (see Figure 4). This interpretation is analogous to the role of *GALNT3*, a Golgi enzyme which acts to O-glycosylate another osteocyte-derived protein, FGF23, protecting it from degradation (51). Given our finding that B4GALNT3 is expressed at high levels in the kidney, a major source of extra-skeletal sclerostin, the kidney may represent the principle source of glycosylated sclerostin. To the extent that activity of renal sclerostin glycosylation influences BMD via changes in circulating sclerostin levels, rather than the latter simply reflecting ‘spill over’ from sclerostin within the bone micro environment, a two-way exchange may exist whereby perturbations in circulating sclerostin levels alter concentrations within bone. The *GALNT1* locus, which was also associated with serum sclerostin, expresses an enzyme in humans that initiates O-glycosylation (52), and may act similarly to alter sclerostin levels by reducing sclerostin clearance.

**Figure 4.**
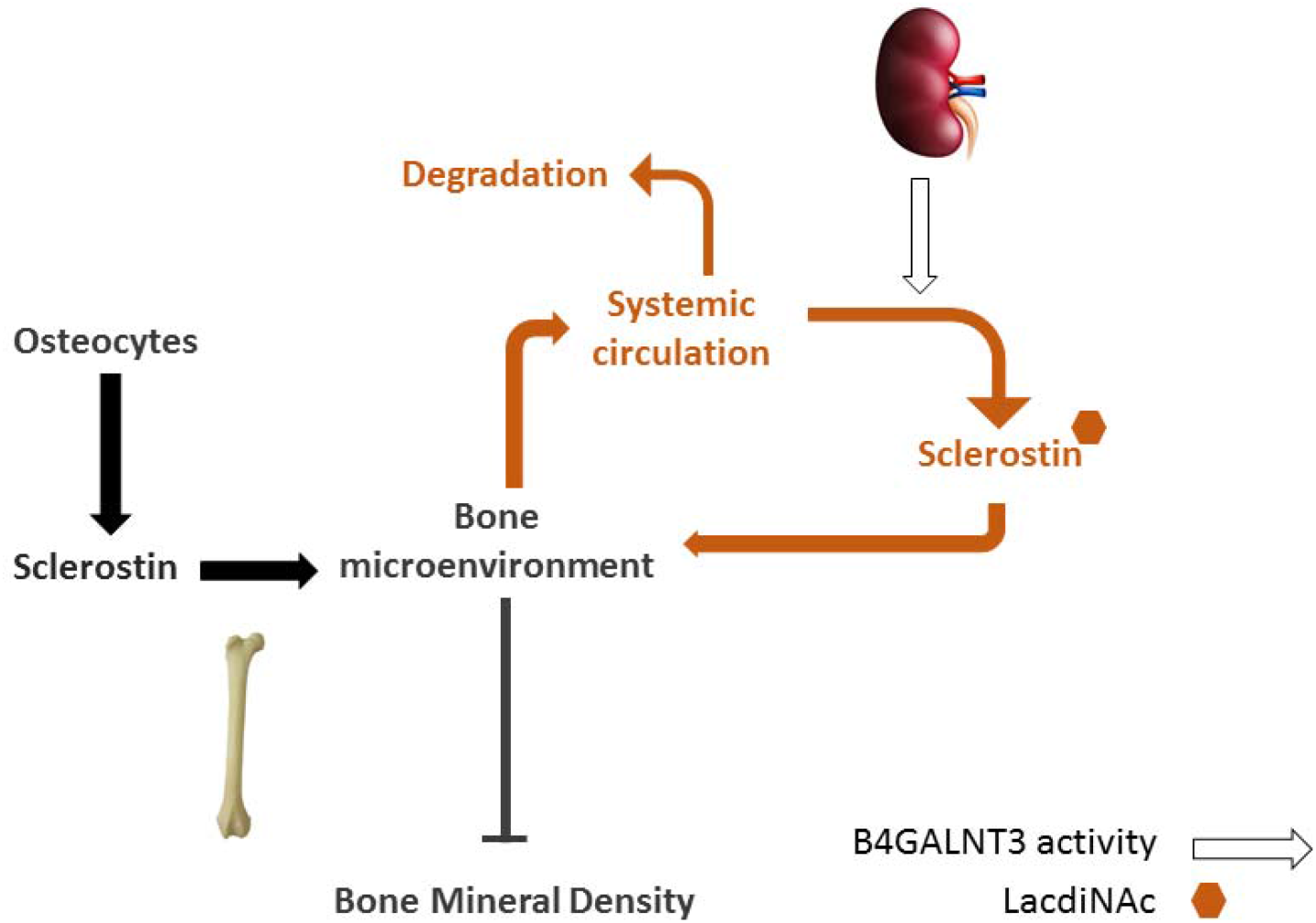
Proposed exchange of sclerostin between skeletal and systemic compartments. Sclerostin, synthesized by osteocytes, is present within the bone micro-environment, but also exchanges with the systemic circulation, where it is produced by several extra-skeletal tissues including the kidney. Whereas locally produced sclerostin is largely responsible for actions of sclerostin on bone, factors which influence circulating levels may also play a role, including variation in activity of B4GALNT3, which we propose protects sclerostin from degradation through generation of a terminal LacdiNAc.

In addition, we found that *B4GALNT3* is expressed at relatively high levels in bone, particularly cortical bone, where sclerostin is also preferentially expressed, reflecting its production by osteocytes. Specifically, *B4GALNT3* mRNA expression was detected in osteoblast cultures derived from mouse calvariae, but not in bone marrow macrophage cultures induced to form osteoclasts. Further studies are required to establish whether B4GALNT3 is expressed in osteocytes as well as osteoblasts. However, to the extent that B4GALNT3 and sclerostin expression overlap within bone, theoretically, B4GALNT3-dependent glycosylation might influence local sclerostin activity in a variety of ways, including altered production. Although genetic variation in *B4GALNT3* might conceivably influence circulating sclerostin via a primary action in bone, against this suggestion, there was little evidence of *B4GALNT3 cis*-regulatory activity in osteoblasts and bone tissue. Furthermore, the *B4GALNT3* SNP did not appear to act as a trans-eQTL signal for *SOST* in bone. However, eQTL studies in bone had considerably less power than those in other tissues analysed (285 and 78 individuals for tibial artery and bone biopsy studies respectively).

Although our analyses were primarily intended to identify loci associated with serum sclerostin, there is some evidence to suggest that rs215224 was responsible for the underlying genetic signal at the *B4GALNT3* locus. This SNP is in perfect LD with our top *B4GALNT3* hit, rs215226. Rs215224 shows strong evidence of alteration of transcriptional activity from RegulomeDB, supported by the finding that this SNP falls within a site of open chromatin, as assessed by ATAC-seq in E15.5 mouse femurs. That said, though rs215224 was also associated with differential methylation at two distinct CPG sites within *B4GALNT3*, one of which coincided with an ATAC-seq site, these were at distinct locations within the gene.

Genetic variation within the *SOST* gene appeared to have relatively weak associations with serum sclerostin, including *SOST* SNPs previously reported to be associated with BMD or hip fracture. This suggests that previously identified *SOST* SNPs influence BMD and hip fracture by altering local sclerostin activity in bone, independently of circulating sclerostin levels. That said, further studies with a larger sample size are needed to establish the contribution of genetic variation within the *SOST* locus to serum sclerostin. In terms of other genetic influences on serum sclerostin, the association between *TNFRSF11B* and serum sclerostin was just below our threshold. *TNFRSF11B* is a well-established BMD locus (42), and the protein product, OPG, plays a major role in regulating bone resorption (53). However, against the suggestion that altered OPG production mediates this genetic association, eQTL studies in heart tissue revealed that rs1485303 (the most strongly associated SNP at this locus) is strongly associated with expression of the adjacent gene, *COLEC10*.

Whereas MR analyses using *B4GLANT3* and *GALNT1* as genetic instruments supported a causal influence of higher sclerostin levels in reducing BMD, bidirectional MR analysis suggested that if anything, higher BMD leads to greater sclerostin levels. This apparent positive influence of BMD on sclerostin levels is in line with previous reports of elevated sclerostin levels in individuals with extreme elevations in BMD (54), and of positive associations between serum sclerostin levels and lumbar spine and femoral neck BMD in postmenopausal women (40). One explanation for this latter relationship is that individuals with higher BMD have a relatively large amount of bone tissue, and therefore, produce more sclerostin as result of having greater numbers of osteocytes. Additionally, these causal relationships may represent components of a regulatory feedback pathway, such that greater bone formation leading to increased BMD stimulates sclerostin production, which then feed backs to reduce bone formation and hence limit BMD gains.

### Strengths and weaknesses

This study represents the first MR study of sclerostin, based on the only serum sclerostin GWAS reported to date. Of the loci identified, *B4GALNT3* had a relatively strong association for a common variant (i.e. 0.2SD change in serum sclerostin per allele). Although we performed a GWAS meta-analysis in order to maximize statistical power, the association between the *B4GALNT3* locus and sclerostin also reached genome wide significance in three of the four participating cohorts. That said, the total GWAS study sample of around 11,000 was relatively small, and further replication of our findings would be desirable. Only two loci passed evidence thresholds for application in our MR analysis, and so it was not possible to apply methods such as Egger regression for evaluating horizontal pleiotropy, to analyse the causal pathway between sclerostin and BMD. Nevertheless, co-localisation analysis strongly suggested that both loci comprising the sclerostin instrument influence BMD via changes in circulating sclerostin, as opposed to a separate, BMD-specific, pathway. Although the genetic association signal between *B4GALNT3* and sclerostin showed high heterogeneity, this appeared to reflect a relatively strong signal in the 4D cohort, comprising individuals with end-stage kidney disease, and is unlikely to limit application as a genetic instrument in the general population. In terms of other limitations, different methods were used to measure sclerostin in the participating cohorts; as well as different ELISAs, the MANOLIS cohort used the OLINK proteomics platform. Nonetheless, similar genetic associations were observed across all four cohorts, suggesting different sclerostin measurement methods are unlikely to have importantly impacted on our results.

### Conclusions

Having applied findings from a GWAS meta-analysis for serum sclerostin in a two sample MR framework, we established a causal relationship between higher circulating sclerostin levels, reduced BMD and increased fracture risk. Hence, strategies for reducing circulating sclerostin may prove valuable in treating osteoporosis. Conceivably, these might include targeting of glycosylation enzymes, encoded by the two genome-wide significant loci for sclerostin we identified.

## ACKNOWLEDGEMENTS

We are extremely grateful to all the families who took part in the ALSPAC study, the midwives for their help in recruiting them, and the whole ALSPAC team, which includes interviewers, computer and laboratory technicians, clerical workers, research scientists, volunteers, managers, receptionists and nurses. ALSPAC data collection was supported by the Wellcome Trust (grants WT092830M; WT088806; WT102215/2/13/2), UK Medical Research Council (G1001357), and University of Bristol. The UK Medical Research Council and the Wellcome Trust (ref: 102215/2/13/2) and the University of Bristol provide core support for ALSPAC. GDS works in the Medical Research Council Integrative Epidemiology Unit at the University of Bristol MC_UU_00011/1.

AG and EZ are supported by Wellcome (098051). CD and CW were supported by the Federal Ministry of Education and Research of the Federal Republic of Germany (BMBF 01EO1504). ML has received lecture or consulting fees from Amgen, Lilly, Meda, UCB Pharma, Renapharma, Radius Health and Consilient Health.

TDC was supported by NSF grant (BCS-1518596). EG was supported by a CIHR Foundation Grant (FDN-148381). KMG and SR were supported by the South East Norway Health Authority under Grant number 52009/8029; the 6th EU framework program under Grant number LSHM-CT-2003-502941; Oslo University Hospital, Ullevaal under Grant number 52009/8029; Lovisenberg Diaconal Hospital. For RNA-Seq analyses we thank Joost Verlouw, Jeroen van Rooij and Masa Zrimšek for their help in the creation, managing and quality control of the RNAseq of data cells derived from iliac crest bone implementing and posterior implementation of the eQTL analysis pipeline. CM-G was supported by the Netherlands Organization for Health Research and Development (ZonMw VIDI 016.136.367).

