## Supplementary documents for "Mendelian Randomization analysis reveals a causal influence of circulating sclerostin levels on bone mineral density and fractures"

Supplementary Results

Supplementary Table 1. GWAS meta-analysis top association signals for circulating sclerostin

See spreadsheet

Supplementary Table 2. The heterogeneity of top association signals across four studies

| SNP (locus) | Beta | SE | P | Study |
| --- | --- | --- | --- | --- |
| rs215226 (*B4GALNT3*) | 0.179 | 0.017 | 1.0E-26 | ALSPAC |
|  | 0.202 | 0.046 | 1.6E-05 | GOOD |
|  | 0.233 | 0.041 | 1.8E-08 | MANOLIS |
|  | 0.344 | 0.043 | 2.2E-15 | 4D |
|  | 0.205 | 0.014 | 4.6E-49 | Overall |
| rs7241221 (*GALNT1*) | 0.105 | 0.020 | 1.1E-07 | ALSPAC |
|  | 0.159 | 0.055 | 3.7E-03 | GOOD |
|  | 0.105 | 0.053 | 4.9E-02 | MANOLIS |
|  | 0.096 | 0.049 | 5.0E-02 | 4D |
|  | 0.109 | 0.017 | 4.4E-11 | Overall |
| rs1485303 (*TNFRSF11B)* | 0.090 | 0.017 | 7.8E-08 | ALSPAC |
|  | 0.000 | 0.044 | 1.0E+00 | GOOD |
|  | 0.102 | 0.040 | 1.2E-02 | MANOLIS |
|  | 0.010 | 0.043 | 8.1E-01 | 4D |
|  | 0.074 | 0.014 | 7.7E-08 | Overall |

Note: locus (nearest gene to the sclerostin associated SNP), Beta (genetic effect per SD change in the minor allele), SE (standard error) and P (p-value for the genetic association).

Supplementary Table 3. The association information of sclerostin SNPs on BMD

| SNP | EA | NONEA | EAF | GENE | BETA_sclerostin | SE_sclerostin | P_sclerostin | BETA_FN-BMD | SE_FN-BMD | P_FN-BMD |
| --- | --- | --- | --- | --- | --- | --- | --- | --- | --- | --- |
| rs215226 | A | G | 0.597 | *B4GALNT3* | 0.205 | 0.014 | 4.60E-49 | -0.017 | 0.008 | 2.81E-02 |
| rs7241221 | G | A | 0.772 | *GALNT1* | 0.109 | 0.017 | 4.40E-11 | -0.002 | 0.009 | 7.93E-01 |
| SNP | EA | NONEA | EAF | GENE | BETA_LS-BMD | SE_LS-BMD | P_LS-BMD | BETA_heel-BMD | SE_heel-BMD | P_heel-BMD |
| rs215226 | A | G | 0.597 | *B4GALNT3* | -0.017 | 0.009 | 6.05E-02 | -0.024 | 0.003 | 5.90E-12 |
| rs7241221 | G | A | 0.772 | *GALNT1* | 0.008 | 0.011 | 4.46E-01 | -0.014 | 0.004 | 1.00E-04 |

Note: EA (Effect allele, orientated in the same direction as the allele related to increasing sclerostin level), NONEA (Other allele), EAF (Effect allele frequency), Beta, SE and P for the listed trait. BMD GWAS lookup (effect sizes, standard errors and P values for the FN-BMD, LS-BMD and eBMD GWAS association separately).

Supplementary Table 4. The association of bone mineral density signals within *SOST* region and their association with sclerostin.

|  |  |  |  |  | BMD / fracture GWAS signals | | | | | | | Sclerostin GWAS lookup | | | | |
| --- | --- | --- | --- | --- | --- | --- | --- | --- | --- | --- | --- | --- | --- | --- | --- | --- |
| SNP | Trait | chrom | BP | Gene | EA | NONEA | | EAF | Beta | SE | P | Proxy_SNP | r^2^ | Beta_ | SE_ | P_ |
| rs7209826 | eBMD | 17 | 41796406 | *SOST* | A | | G | 0.620 | -0.042 | 0.004 | 2E-36 | rs12600549 | 0.703 | 0.028 | 0.014 | 0.049 |
| rs188810925 | eBMD | 17 | 41798194 | *SOST* | G | | A | 0.932 | -0.067 | 0.007 | 1E-26 | rs66838809 | 0.888 | 0.058 | 0.027 | 0.032 |
| rs170634 | eBMD | 17 | 42175821 | *SOST* | A | | C | 0.744 | -0.031 | 0.004 | 6E-17 | rs850856 | 0.782 | 0.016 | 0.017 | 0.334 |
| rs1513670 | FN-BMD | 17 | 41807331 | *SOST* | T | | C | 0.366 | -0.027 | 0.004 | 3E-16 | rs9303540 | 0.983 | 0.026 | 0.014 | 0.070 |
| rs2741856 | fracture | 17 | 41826839 | *SOST* | G | | C | 0.920 | 0.095 | 0.014 | 3E-25 | rs66838809 | 0.651 | 0.058 | 0.027 | 0.032 |

Note: Trait [BMD estimated from heel ultrasound (eBMD), femoral neck BMD (FN-BMD)], EA (Effect allele, orientated in the same direction as the (proxy SNP) allele related to increasing sclerostin level), NONEA (Other allele), EAF (Effect allele frequency), Beta, SE and P for the listed trait. Proxy_SNP is the proxy sclerostin SNP with strongest linkage disequilibrium with the BMD/fracture GWAS SNP; r2 (pairwise linkage disequilibrium between the BMD/fracture SNP and the proxy sclerostin SNP); Sclerostin GWAS lookup (effect sizes, standard errors and P values for the sclerostin GWAS association).

Supplementary Table 5. The genetic correlation between sclerostin and 11 bone phenotypes in LD Hub

| trait1 | trait2 | Genetic correlation | | | | | | SNP heritability | | | |
| --- | --- | --- | --- | --- | --- | --- | --- | --- | --- | --- | --- |
|  |  | r_g_ | se | z | p | gcov_int | gcov_int_se | h2_obs | h2_obs_se | h2_int | h2_int_se |
| Sclerostin | Fracture by fall | -0.246 | 0.306 | -0.804 | 0.421 | 0.001 | 0.005 | 0.036 | 0.014 | 0.995 | 0.008 |
| Sclerostin | Osteoporosis | -0.198 | 0.204 | -0.972 | 0.331 | -0.010 | 0.005 | 0.009 | 0.002 | 1.021 | 0.009 |
| Sclerostin | Fracture of forearm | -0.072 | 0.256 | -0.283 | 0.778 | 0.002 | 0.006 | 0.004 | 0.002 | 1.010 | 0.008 |
| Sclerostin | eBMD | 0.068 | 0.100 | 0.676 | 0.499 | 0.010 | 0.009 | 0.238 | 0.036 | 1.210 | 0.048 |
| Sclerostin | eBMD (left) | 0.096 | 0.114 | 0.839 | 0.401 | 0.007 | 0.007 | 0.230 | 0.036 | 1.111 | 0.029 |
| Sclerostin | eBMD (right) | 0.173 | 0.117 | 1.476 | 0.140 | 0.001 | 0.007 | 0.235 | 0.037 | 1.108 | 0.029 |
| Sclerostin | FN-BMD 2012 | 0.182 | 0.146 | 1.249 | 0.212 | 0.022 | 0.006 | 0.199 | 0.022 | 1.019 | 0.010 |
| Sclerostin | LS-BMD 2012 | 0.283 | 0.203 | 1.39 | 0.165 | 0.012 | 0.005 | 0.070 | 0.014 | 1.009 | 0.010 |
| Sclerostin | FN-BMD 2015 | 0.325 | 0.187 | 1.735 | 0.083 | 0.022 | 0.005 | 0.070 | 0.012 | 1.006 | 0.009 |
| Sclerostin | LS-BMD 2015 | 0.359 | 0.179 | 2.007 | 0.045 | 0.031 | 0.006 | 0.164 | 0.021 | 1.051 | 0.011 |
| Sclerostin | FA-BMD 2015 | 0.551 | 0.761 | 0.724 | 0.469 | -0.006 | 0.005 | 0.026 | 0.038 | 1.026 | 0.007 |

Note: Genetic correlation between sclerostin and bone phenotypes (genetic correlations (r_g_), standard error (se), P value (p), intercept (gcov_int), standard error of intercept (gcov_int_se)); SNP heritability for the bone phenotypes (observed SNP heritability (h^2^), standard error (se), intercept (h2_int) and standard error of intercept (h2_int_se)). Abbreviations: eBMD (bone mineral density estimated by heel ultrasound), LS-BMD (lumbar spine bone mineral density), FN-BMD (femoral Neck bone mineral density), FA-BMD (Forearm Bone mineral density), 2012 (BMD paper published by Estrada et al in 2012) and 2015 (BMD paper published by Zheng et al 2015).

Supplementary Table 6. The Mendelian randomization results for sclerostin on bone phenotypes and the heterogeneity test of genetic instruments

| Outcome | n | SNP | | b | se | p | Q | Q_df | Q_pval |
| --- | --- | --- | --- | --- | --- | --- | --- | --- | --- |
| Femoral neck bone mineral density | 32744 | rs215226 | -0.119 | | 0.040 | 0.0030 | NR | NR | NR |
| Femoral neck bone mineral density | 32744 | rs7241221 | -0.147 | | 0.090 | 0.100 | NR | NR | NR |
| Femoral neck bone mineral density | 32744 | IVW | -0.123 | | 0.037 | 0.00074 | 0.086 | 1 | 0.770 |
| Lumbar spine bone mineral density | 31800 | rs215226 | -0.155 | | 0.042 | 0.00026 | NR | NR | NR |
| Lumbar spine bone mineral density | 31800 | rs7241221 | 0.073 | | 0.095 | 0.442 | NR | NR | NR |
| Lumbar spine bone mineral density | 31800 | IVW | -0.117 | | 0.085 | 0.167 | 4.802 | 1 | 0.028 |
| eBMD_500K | 426824 | rs215226 | -0.118 | | 0.009 | 1.97E-37 | NR | NR | NR |
| eBMD_500K | 426824 | rs7241221 | -0.143 | | 0.021 | 3.08E-12 | NR | NR | NR |
| eBMD_500K | 426824 | IVW | -0.122 | | 0.009 | 1.29E-38 | 1.248 | 1 | 0.264 |
| Fracture_500K | 426795 | rs215226 | 0.087 | | 0.029 | 0.0029 | NR | NR | NR |
| Fracture_500K | 426795 | rs7241221 | 0.227 | | 0.065 | 0.00046 | NR | NR | NR |
| Fracture_500K | 426795 | IVW | 0.111 | | 0.052 | 0.034 | 3.851 | 1 | 0.050 |

Note: SNP, the rsid of the genetic instruments; b is the causal effect of sclerostin on bone phenotypes; se and p are the standard error, upper 95% confidence interval, lower 95% confidence internal and P value of the causal estimates. N is the sample size of the outcome GWAS. Q, Q_df and Q_pval are the Cochrane Q value, degree of freedom and P value of the heterogeneity test of the genetic instruments. Abbreviations: eBMD_500K (bone mineral density estimated by heel ultrasound, published by Morris et al 2018), Fracture_500K (published by Morris et al 2018), LS-BMD (lumbar spine bone mineral density, published by Estrada et al in 2012), FN-BMD (femoral Neck bone mineral density, published by Estrada et al in 2012), IVW (Inverse variance weighted)

Supplementary Table 7. Colocalization analysis results for genetic associations with sclerostin and bone phenotypes

| Outcome | n | SNP | nsnps-coloc | | PP.H0 | PP.H1 | PP.H2 | PP.H3 | PP.H4 |
| --- | --- | --- | --- | --- | --- | --- | --- | --- | --- |
| FN-BMD | 49988 | rs215226 | | 2221 | 7.45E-21 | 8.91E-01 | 4.59E-22 | 5.49E-02 | 5.37E-02 |
| FN-BMD | 49988 | rs7241221 | | 2065 | 4.15E-02 | 9.10E-01 | 1.82E-03 | 3.99E-02 | 6.71E-03 |
| LS-BMD | 44731 | rs215226 | | 2220 | 7.67E-21 | 9.18E-01 | 3.89E-22 | 4.66E-02 | 3.57E-02 |
| LS-BMD | 44731 | rs7241221 | | 2065 | 3.94E-02 | 8.65E-01 | 3.83E-03 | 8.40E-02 | 7.63E-03 |
| eBMD_500K | 426824 | rs215226 | | 2386 | 2.69E-53 | 3.22E-33 | 3.18E-23 | 2.80E-03 | 9.97E-01 |
| eBMD_500K | 426824 | rs7241221 | | 2152 | 6.33E-10 | 1.41E-08 | 1.32E-04 | 1.95E-03 | 9.98E-01 |
| Fracture500K | 426795 | rs215226 | | 2477 | 6.56E-21 | 7.85E-01 | 2.46E-22 | 2.92E-02 | 1.86E-01 |
| Fracture500K | 426795 | rs7241221 | | 2192 | 1.72E-02 | 3.83E-01 | 4.58E-03 | 1.02E-01 | 4.94E-01 |

Note: nsnps-coloc, the number of SNPs included in the colocalization analysis; PP.H0 to PP.H4 are the posterior probability (PP) of each colocalization hypothesis tested: H0 (neither trait associated), H1 (sclerostin associated only), H2 (outcome associated only), H3 (both sclerostin and outcome associated, but with different causal variants), H4 (colocalization, both traits associated with the same causal variant). Abbreviations: eBMD_500K (bone mineral density estimated by heel ultrasound, published by Morris et al 2018), Fracture500K (published by Morris et al 2018), LS-BMD (lumbar spine bone mineral density), FN-BMD (femoral Neck bone mineral density, published by Estrada et al in 2012), LS-BMD (lumbar spine Bone mineral density, published by Estrada et al in 2012).

Supplementary Table 8. Phenome-wide association study results for top sclerostin associated SNP -- rs215226.

See spreadsheet

Supplementary Table 9. Phenome-wide association study results for top sclerostin associated SNP -- rs7241221.

See spreadsheet

Supplementary Table 10. Observational correlation between standing height and sitting hit on BMD

| Trait1 | ID_Trait1 | Trait2 | ID_Trait2 | Correlation | P-val |
| --- | --- | --- | --- | --- | --- |
| Standing height | 50 | Heel bone mineral density (BMD) | 3148 | 0.10 | 0 |
| Sitting height | 20015 | Heel bone mineral density (BMD) | 3148 | 0.11 | 0 |
| Standing height | 50 | Heel bone mineral density (BMD) T-score, automated | 78 | 0.10 | 0 |
| Sitting height | 20015 | Heel bone mineral density (BMD) T-score, automated | 78 | 0.11 | 0 |

Note: ID is the UK Biobank ID for the trait, correlation is the Pearson correlation between each pair of traits, P-val is the P value of the correlation analysis. The data is from UK Biobank.

Supplementary Table 11. The genetic association information of three top sclerostin association signals adjust for human height using data from ALSPAC cohort.

See spreadsheet

Supplementary Table 7. Instruments of the bidirectional MR of femoral neck and lumbar spine bone mineral density on sclerostin and Steiger filtering results of the BMD instruments.

See spreadsheet

Supplementary Table 13. Bidirectional MR results using femoral neck and lumbar spine bone mineral density as exposure and sclerostin as outcome.

| exposure | method | nsnp | b | se | LCI | UCI | P-val |
| --- | --- | --- | --- | --- | --- | --- | --- |
| FN-BMD | MR Egger | 33 | 0.320 | 0.178 | -0.029 | 0.670 | 0.082 |
| FN-BMD | Weighted median | 33 | 0.173 | 0.077 | 0.021 | 0.324 | 0.026 |
| FN-BMD | Inverse variance weighted | 33 | 0.167 | 0.060 | 0.050 | 0.285 | 0.005 |
| LS-BMD | MR Egger | 35 | 0.369 | 0.201 | -0.024 | 0.763 | 0.075 |
| LS-BMD | Weighted median | 35 | 0.213 | 0.073 | 0.069 | 0.356 | 0.004 |
| LS-BMD | Inverse variance weighted | 35 | 0.195 | 0.059 | 0.080 | 0.310 | 0.0009 |
| FN-BMD | MR Egger (no outlier SNP) | 30 | 0.287 | 0.152 | -0.012 | 0.585 | 0.070 |
| FN-BMD | Weighted median (no outlier SNP) | 30 | 0.134 | 0.075 | -0.013 | 0.281 | 0.074 |
| FN-BMD | Inverse variance weighted (no outlier SNP) | 30 | 0.149 | 0.053 | 0.045 | 0.253 | 0.005 |
| LS-BMD | MR Egger (no outlier SNP) | 32 | 0.435 | 0.173 | 0.096 | 0.773 | 0.017 |
| LS-BMD | Weighted median (no outlier SNP) | 32 | 0.233 | 0.075 | 0.085 | 0.381 | 0.002 |
| LS-BMD | Inverse variance weighted (no outlier SNP) | 32 | 0.229 | 0.053 | 0.126 | 0.332 | 1.4x10^-05^ |

Note: b is the causal estimate of femoral neck bone mineral density on sclerostin. se, LCI, UCI and P-val are the standard error, upper 95% confidence interval, lower 95% confidence internal and P value of the causal estimates. Number of SNPs in the genetic instrument was 32. Abbreviations: FN-BMD (femoral Neck bone mineral density, published by Estrada et al in 2012); LS-BMD (lumbar spine Bone mineral density, published by Estrada et al in 2012). No outlier SNP (the MR using BMD associated SNPs excluding outliers SNPs using Radial MR approach)

Supplementary Table 14. Regulomedb score for top association signals of sclerostin

| Leading_SNP | Proxy_SNP | Rseq | Locus | RegulomeDB_score | |
| --- | --- | --- | --- | --- | --- |
| rs215226 | rs4980826 | 0.850 | B4GALNT3 | | 2b |
| rs215226 | rs6489548 | 0.984 | B4GALNT3 | | 6 |
| rs215226 | rs7294354 | 0.867 | B4GALNT3 | | 6 |
| rs215226 | rs56279417 | 0.859 | B4GALNT3 | | 5 |
| rs215226 | rs215223 | 0.996 | B4GALNT3 | | 4 |
| rs215226 | rs215224 | 1.000 | B4GALNT3 | | 2b |
| rs215226 | rs215225 | 0.871 | B4GALNT3 | | 4 |
| rs215226 | rs215226 | 1.000 | B4GALNT3 | | 5 |
| rs7241221 | rs7241221 | 1.000 | GALNT1  GALNT1 | | 5 |
| rs7241221 | rs8095921 | 0.970 |  |  | 6 |
| rs1485303 | rs3134054 | 0.817 | TNFRSF11B | | 6 |
| rs1485303 | rs11573885 | 0.862 | TNFRSF11B | | 5 |
| rs1485303 | rs1872426 | 0.866 | TNFRSF11B | | 1f |
| rs1485303 | rs6993910 | 0.866 | TNFRSF11B | | 7 |
| rs1485303 | rs1905786 | 0.866 | TNFRSF11B | | 7 |
| rs1485303 | rs3134056 | 0.817 | TNFRSF11B | | 7 |
| rs1485303 | rs3134057 | 0.821 | TNFRSF11B | | 4 |
| rs1485303 | rs1485289 | 0.862 | TNFRSF11B | | 7 |
| rs1485303 | rs7464496 | 0.862 | TNFRSF11B | | 6 |
| rs1485303 | rs2073618 | 0.875 | TNFRSF11B | | 4 |
| rs1485303 | rs1564861 | 0.827 | TNFRSF11B | | 6 |
| rs1485303 | rs1485302 | 1.000 | TNFRSF11B | | 5 |
| rs1485303 | rs1385500 | 1.000 | TNFRSF11B | 5 | |
| rs1485303 | rs1485303 | 1.000 | TNFRSF11B | 5 | |

Note: Leading SNP is the top association signal of sclerostin within each sclerostin loci; Proxy SNP is the SNP with r^2^>0.8 with the leading SNP; Rseq is the LD r^2^ between leading SNP and proxy SNP; Gene is the nearby gene of the leading SNP; RegulomeDB_score is the functional score of the proxy SNP from RegulomeDB database.

Supplementary Table 15. ATAC-seq results for the top association signals of sclerostin

| Chr | POS | SNP | Tissue source for ATAC-seq |
| --- | --- | --- | --- |
| chr8 | 119964052 | rs2073618 | Proximal femur |
| chr12 | 590259 | rs215223 | Proximal femur |
| chr12 | 590449 | rs215224 | Proximal femur |
| chr12 | 590545 | rs215225 | Proximal femur |
| chr8 | 119964052 | rs2073618 | Distal femur |
| chr12 | 589813 | rs12318530 | Distal femur |
| chr12 | 590259 | rs215223 | Distal femur |
| chr12 | 590449 | rs215224 | Distal femur |
| chr12 | 590545 | rs215225 | Distal femur |

Note: SNP represents the sclerostin-associated SNPs overlapping with ATAC-seq peaks.

Supplementary Table 16. Cis eQTL lookup for the top sclerostin association signals in multiple tissues from GTEx consortium

| Gene Symbol | Effect allele / Other allele | SNP | P-Value | Beta | Tissue | |
| --- | --- | --- | --- | --- | --- | --- |
| B4GALNT3 | A / G | rs215226 | 6.70E-66 | 0.640 | | Artery - Tibial |
| B4GALNT3 | A / G | rs215226 | 1.30E-30 | 0.510 | | Artery - Aorta |
| B4GALNT3 | A / G | rs215226 | 1.90E-12 | 0.510 | | Artery - Coronary |
| B4GALNT3 | A / G | rs215226 | 3.70E-07 | 0.560 | | Ovary |
| B4GALNT3 | A / G | rs215226 | 1.30E-06 | -0.150 | | Nerve - Tibial |
| INO80C | G / A | rs7241221 | 1.10E-06 | 0.220 | | Muscle - Skeletal |
| GALNT1 | G / A | rs7241221 | 1.40E-05 | 0.190 | | Adipose - Subcutaneous |
| INO80C | G / A | rs7241221 | 1.70E-05 | 0.240 | | Adipose - Subcutaneous |
| COLEC10 | G / A | rs1485303 | 5.30E-12 | 0.400 | | Heart - Left Ventricle |

Note: Gene Symbol is the ENSG id and gene symbol of the gene expression probe; Variant Id and SNP are the cptid and rsid of the eQTL; beta is the effect size of the eQTL on the expression.

Supplementary Table 17. Colocalization analysis of eQTL and circulating sclerostin within the sclerostin loci

| Tissue | PP.H0.abf | PP.H1.abf | PP.H2.abf | PP.H3.abf | PP.H4.abf |
| --- | --- | --- | --- | --- | --- |
| Artery Aorta | 8.79E-56 | 1.02E-35 | 5.50E-23 | 5.39E-03 | 0.995 |
| Artery Coronary | 1.06E-51 | 1.23E-31 | 5.50E-23 | 5.40E-03 | 0.995 |
| Artery Tibial | 1.96E-121 | 2.27E-101 | 5.49E-23 | 5.38E-03 | 0.995 |
| Ovary | 6.72E-24 | 7.80E-04 | 5.68E-23 | 5.60E-03 | 0.994 |
| Nerve Tibial | 9.70E-30 | 1.13E-09 | 8.61E-21 | 1.000 | 6.31E-07 |

Note: PP.H0.abf to PP.H4.abf are the probability of each colocalization hypothesis.

Supplementary Table 18. Cis eQTL lookup for the top sclerostin association signals in osteoblast cell

See spreadsheet

Supplementary Table 19. Cis eQTL lookup for the top sclerostin association signals in iliac crest bone biopsies

| Gene | HGNCName | ProbeChr | ProbeCenterPos | SNP-ID | SNPChr | SNPPos | OA/EA | Beta | SE | P | Sample_size | P_sclerostin |
| --- | --- | --- | --- | --- | --- | --- | --- | --- | --- | --- | --- | --- |
| *B4GALNT3* | ENSG00000139044.6 | 12 | 621102 | rs215226 | 12 | 591300 | A/G | 0.012 | 0.115 | 0.917 | 78 | 4.6E-49 |
| *GALNT1* | ENSG00000141429.9 | 18 | 33226439 | rs7241221 | 18 | 33152792 | G/A | 0.011 | 0.115 | 0.924 | 78 | 4.4E-11 |
| *TNFRSF11B* | ENSG00000164761.4 | 8 | 119950117 | rs1485303 | 8 | 119976256 | G/A | -0.199 | 0.112 | 0.081 | 78 | 7.7E-08 |

Note: Gene information of the probe (Gene (gene symbol), NGNCName (ENSG id of the gene), ProbeChr (the chromosome of the probe), ProbeCenterPos (the center position of the probe)). eQTL information (SNP ID (the rsid of the eQTL), SNPChr (the chromosome of the eQTL), SNPPos (the position of the eQTL), OA/EA (the effect allele and other allele of the pQTL), Beta (the effect size of the association between SNP and the gene expression level of the probe), SE (standard error of the association), P (P value of the association), p_sclerostin (P value of the association between the SNP and circulating sclerostin level)

Supplementary Table 20. Trans eQTL lookup for the top sclerostin association signals within SOST region in iliac crest bone biopsies

See spreadsheet

Supplementary Table 21. Cis methylation QTL lookup for the top sclerostin association signals in blood usig ALSPAC methylation data (ARIES)

| Timepoint | SNP | Nearby_gene | Chr | Pos | A1 | A2 | MAF | CpG | CpG Chr | CpG Pos | CpG-island-start | | CpG-island-end | | Beta | P-value |
| --- | --- | --- | --- | --- | --- | --- | --- | --- | --- | --- | --- | --- | --- | --- | --- | --- |
| Adolescence | rs1485303 | TNFRSF11B | 8 | 119976256 | A | G | 0.438 | cg13268132 | 8 | 119964037 | | 119963946 | | 119964178 | 0.394 | 3.96E-20 |
| Adolescence | rs1485303 | TNFRSF11B | 8 | 119976256 | A | G | 0.438 | cg17171407 | 8 | 119960777 | | NA | | NA | 0.478 | 6.86E-37 |
| Birth | rs1485303 | TNFRSF11B | 8 | 119976256 | A | G | 0.443 | cg13268132 | 8 | 119964037 | | 119963946 | | 119964178 | 0.387 | 1.35E-18 |
| Birth | rs1485303 | TNFRSF11B | 8 | 119976256 | A | G | 0.443 | cg17171407 | 8 | 119960777 | | NA | | NA | 0.249 | 2.45E-11 |
| Childhood | rs1485303 | TNFRSF11B | 8 | 119976256 | A | G | 0.438 | cg13268132 | 8 | 119964037 | | 119963946 | | 119964178 | 0.473 | 6.75E-32 |
| Childhood | rs1485303 | TNFRSF11B | 8 | 119976256 | A | G | 0.438 | cg17171407 | 8 | 119960777 | | NA | | NA | 0.509 | 2.54E-45 |
| Middle Age | rs1485303 | TNFRSF11B | 8 | 119976256 | A | G | 0.453 | cg13268132 | 8 | 119964037 | | 119963946 | | 119964178 | 0.441 | 2.75E-24 |
| Middle Age | rs1485303 | TNFRSF11B | 8 | 119976256 | A | G | 0.453 | cg17171407 | 8 | 119960777 | | NA | | NA | 0.504 | 2.50E-42 |
| Pregnancy | rs1485303 | TNFRSF11B | 8 | 119976256 | A | G | 0.455 | cg17171407 | 8 | 119960777 | | NA | | NA | 0.551 | 1.48E-45 |
| Pregnancy | rs1485303 | TNFRSF11B | 8 | 119976256 | A | G | 0.455 | cg13268132 | 8 | 119964037 | | 119963946 | | 119964178 | 0.442 | 4.88E-27 |
| Adolescence | rs215226 | B4GALNT3 | 12 | 591300 | G | A | 0.409 | cg20907806 | 12 | 570249 | | 568702 | | 570362 | -0.264 | 1.13E-11 |
| Childhood | rs215226 | B4GALNT3 | 12 | 591300 | G | A | 0.412 | cg20907806 | 12 | 570249 | | 568702 | | 570362 | -0.263 | 8.19E-11 |
| Middle Age | rs215226 | B4GALNT3 | 12 | 591300 | G | A | 0.391 | cg20907806 | 12 | 570249 | | 568702 | | 570362 | -0.286 | 6.70E-10 |
| Pregnancy | rs215226 | B4GALNT3 | 12 | 591300 | G | A | 0.389 | cg20907806 | 12 | 570249 | | 568702 | | 570362 | -0.254 | 1.02E-08 |
| Pregnancy | rs215226 | B4GALNT3 | 12 | 591300 | G | A | 0.389 | cg26388816 | 12 | 570155 | | 568702 | | 570362 | -0.262 | 1.85E-09 |

Note: Timepoint, the time point the DNA methylation were measured. mQTL information (SNP (the rsid of the mQTL), Nearby_gene (the gene near the mQTL), Chr (the chromosome of the mQTL), Pos (the position of the mQTL), A1 and A2 (the effect allele and other allele of the mQTL), MAF (minor allele frequency of the mQTL), Beta (the effect size of the association between SNP and the DNA methylation level of the probe), P-value (P value of the association)). DNA methylation level of the probe (CpG (the CpG id of the probe), CpG Chr (the chromosome of the probe), CpG Pos(the position of the probe), CpG-island-start and CpG-island-end (the start and end position of the CpG island)).

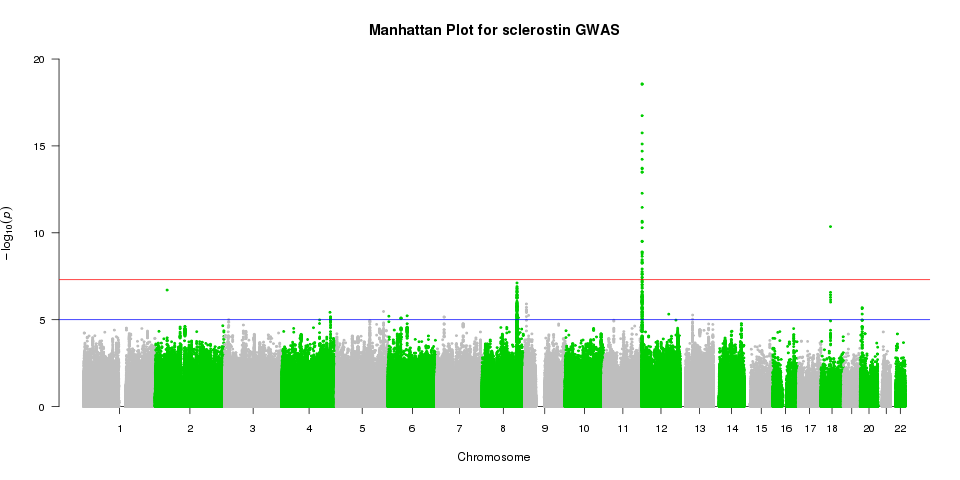

Supplementary Figure 1. Manhattan plot of the sclerostin GWAS meta-analysis. The X-axis indicates the chromosomal position of each SNP, whereas the Y-axis denotes the evidence of association shown as −log(P-value). The red line indicates genome-wide significance of association (P = 5×10^−8^), the blue line indicates suggestive association (P = 1×10^−5^).

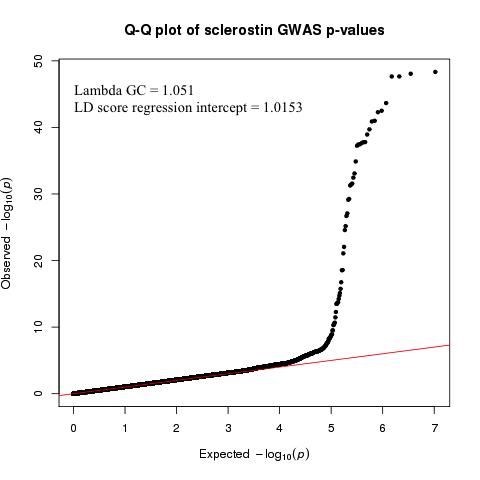

Supplementary Figure 2. QQ plots from the GWAS meta-analysis of sclerostin. The Lambda of genomic inflation factor is equal to 1.048. The LD score regression intercept is equal to 1.0007.

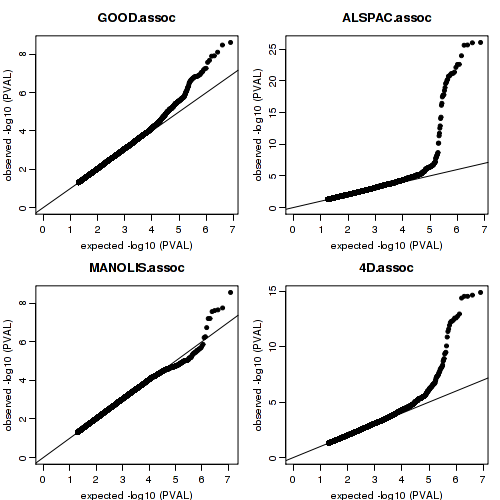

Supplementary Figure 3. QQ plot for GWASs using ALSPCA, 4D, GOOD and MANOLIS data.

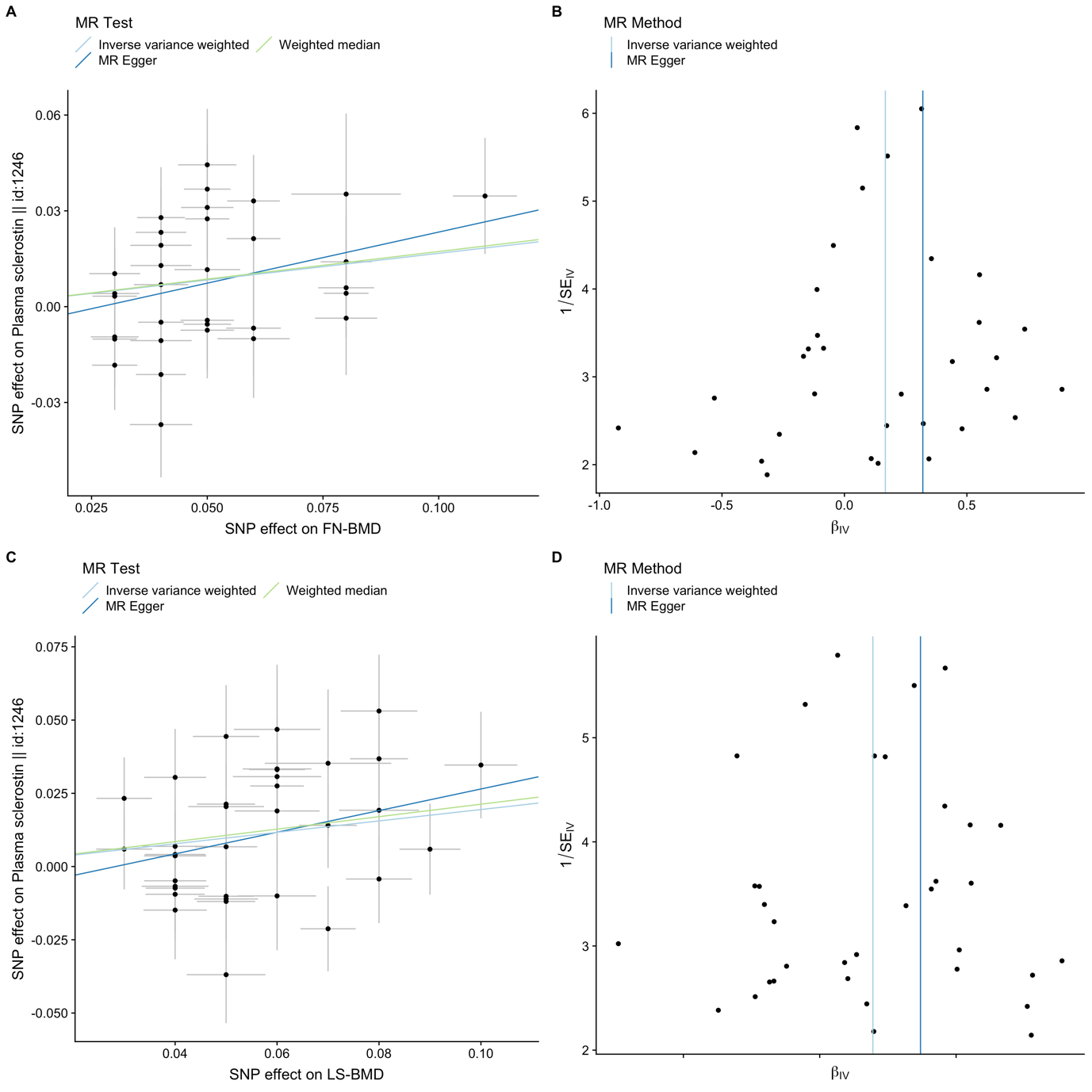

Supplementary Figure 4. Scatter plots displaying estimates of the association between each SNP and sclerostin (y-axis) against estimates of the association between each SNP and BMD level (x-axis), i.e femoral neck BMD (panel A) and lumbar spine BMD (panel C); and funnel plots displaying instrument strength (y-axis) plotted against causal effect estimate (x-axis) for 33 SNPs associated with femoral neck BMD (Panel B) and 35 SNPs associated with lumbar spine BMD (Panel D). The inverse variance weighted, MR Egger and weighted median causal effect estimates are represented by a light blue, dark blue and green respectively. The y-intercept of the dark blue regression line denotes the estimate of the degree of directional pleiotropy in the dataset. The inverse-variance weighted causal effect estimate is represented by the slope of the light blue line.

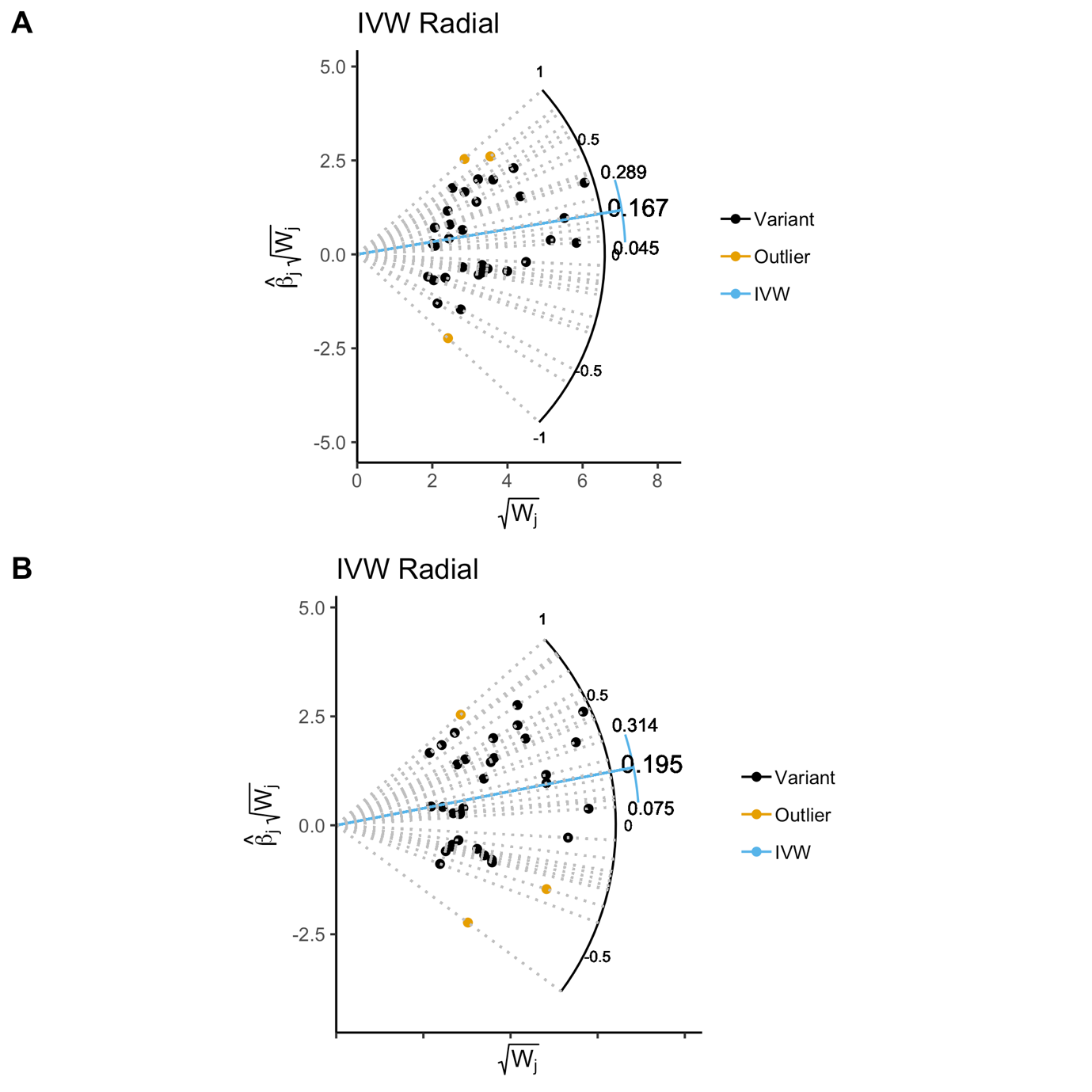

Supplementary Figure 5. Radial plot for detecting outlier instruments for A) FN-BMD; and B) LS-BMD. The line in blue is the regression line of the IVW approach. Outlier SNP is in yellow and other variants in black.

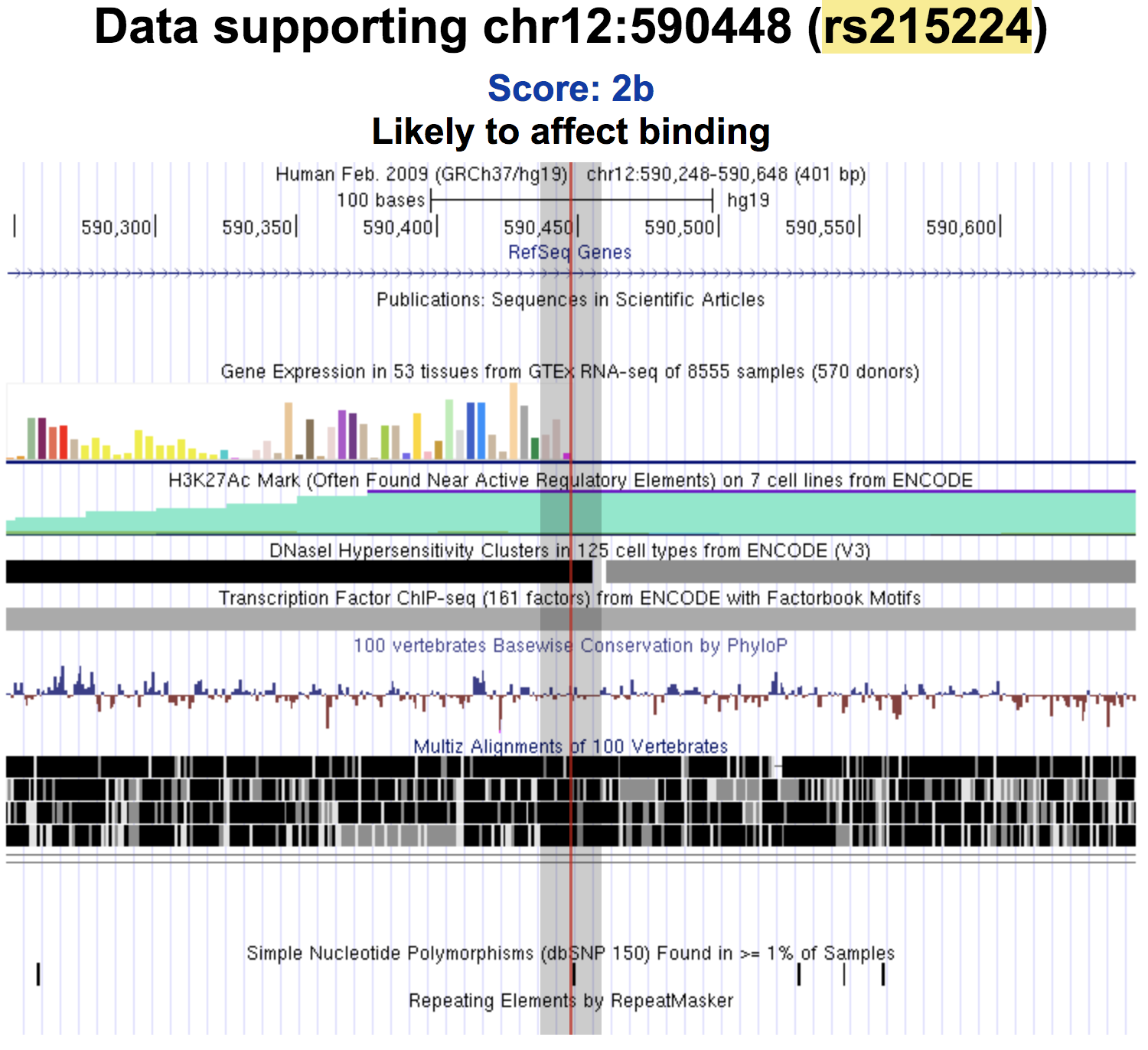

Supplementary Figure 6. Regulomedb results for the top association signal in *B4GLANT3*.

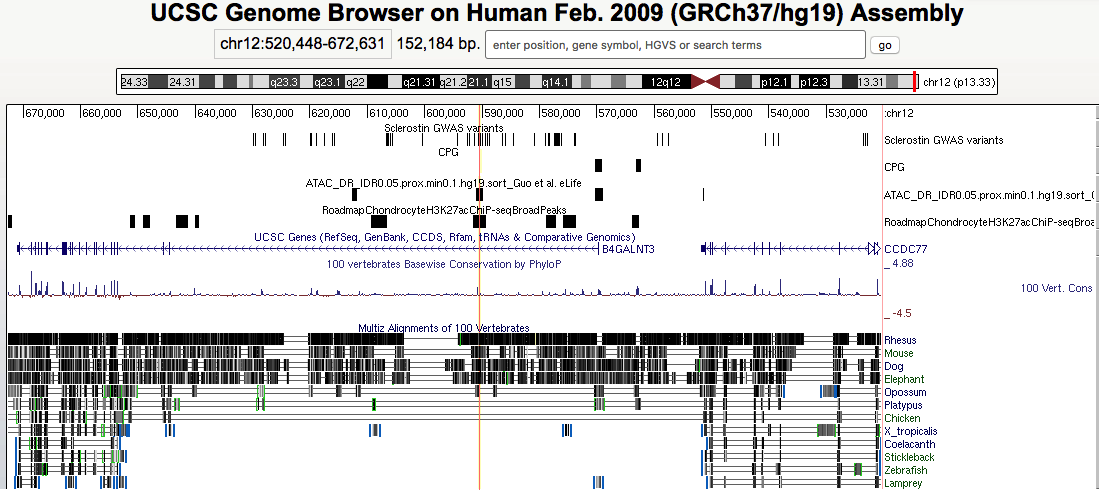

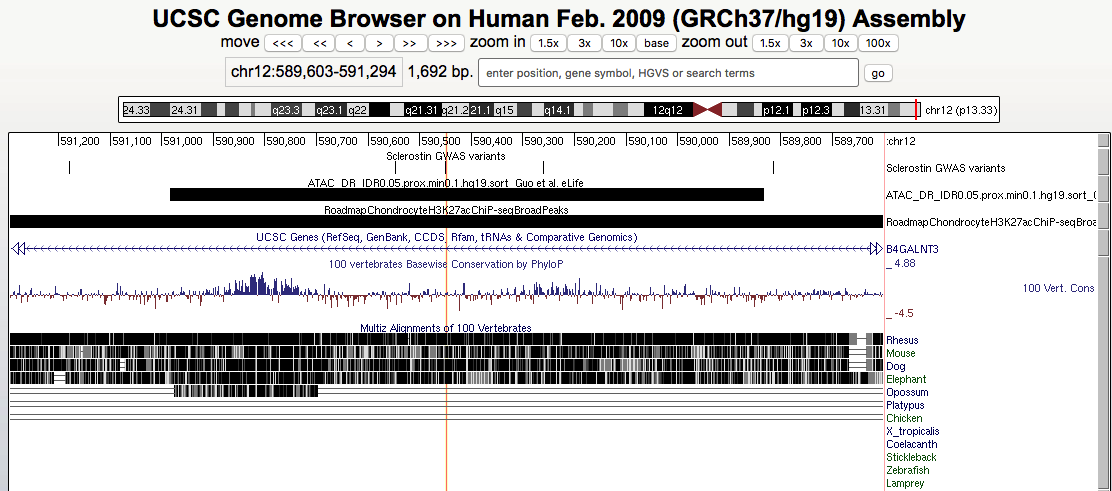

rs215224

rs215224

Supplementary Figure 7. ATAC-seq results within the *B4GALNT3* locus. Upper panel: the zoom-out plot showing the entire *B4GLANT3* locus. Lower panel: zoom-in view of the genomic region containing 4 sclerostin associated SNPs within the *B4GALNT3* region. The red line is the location of the potential functional SNP in the *B4GALNT3* region – rs215224. This SNP overlapped with an ATAC-seq peak as well as a ChiIP-seq peak.
